# Hierarchical Cognition causes Task Related Deactivations but not just in Default Mode Regions

**DOI:** 10.1101/297259

**Authors:** Ausaf A Farooqui, Tom Manly

**Affiliations:** MRC-Cognition & Brain Sciences Unit, Cambridge, UK

**Author notes:** Correspondence: Ausaf A Farooqui, Medical Research Council Cognition & Brain Sciences Unit, Cambridge, UK, CB2 7EF.

## Abstract

The well-known deactivation of the Default Mode Network (DMN) during external tasks is usually thought to reflect the suppression of internally directed mental activity during external attention. In 3 experiments with human participants we organized sequences of task events identical in their attentional and control demands into larger task episodes. We found that DMN deactivation across such sequential events was never constant, but was maximum at the beginning of the episode, then decreased gradually across the episode, reaching baseline towards episode completion, with the final event of the episode eliciting an activation. Crucially, this pattern of activity was not limited to a fixed set of DMN regions but, across experiments, was shown by a variable set of regions expected to be uninvolved in processing the ongoing task. This change in deactivation across sequential but identical events showed that the deactivation cannot be related to attentional/control demands which were constant across the episode, instead it has to be related to some episode related load that was maximal at the beginning and then decreased gradually as parts of the episode got executed. We argue that this load resulted from cognitive programs through which the entire episode was hierarchically executed as one unit. At the beginning of task episodes, programs related to their entire duration is assembled, causing maximal deactivation. As execution proceeds, elements within the program related to the completed parts of the episode dismantle, thereby decreasing the program load and causing a decrease in deactivation.

**Significance Statement:** We prepare breakfasts and write emails, and not individually execute their many component acts. The current study suggests that cognitive entities that enable such hierarchical execution of goal-directed behavior may cause the ubiquitously seen deactivation during external task execution. Further, while this deactivation has previously been associated with a defined set of the so called default mode regions, current study demonstrates that deactivation is shown by *any* region not *currently* involved in task execution, and in certain task episodes can even include attention related fronto-parietal regions as well as primary sensory and motor regions.

## Introduction

Default mode network (DMN) regions are well known to *deactivate* during external task execution (Raichle et al., 2001). This deactivation has been attributed to their being specialized for internally directed cognitive activities like mind wandering, theory of mind and autobiographical memories, which causes them to deactivate when attention and control is directed to external stimuli (Buckner et al., 2008; Spreng et al., 2009). These regions are contrasted with Multiple Demands (MD) regions, a set of fronto-parietal regions that are thought to always *activate* during external task executions (Duncan, 2013). Here we show that task related deactivation is primarily related to cognitive entities through which task episodes are executed as one unit, not to external attention or control per se.

Attention and control are always instantiated in the context of task episodes - extended periods during which cognition is focused on a particular goal and generates a sequence of actions to complete it e.g. preparing breakfast, writing email, a run of trials in experimental sessions etc. Task episodes, despite being temporally extended and consisting of a sequence of smaller acts, are executed as one unit, and not as a collection of independent acts, because their execution occurs through *programs* that are instantiated as one entity but contain elements related to the entire episode (see also plans, scripts and schemas; Miller et al., 1960; Schank and Abelson, 1977; Cooper and Shallice, 2000). Such programs are assembled at the beginning of task episode execution and are evidenced by the slower step 1 RTs of task episodes (Schneider and Logan, 2006), and by the fact that this step 1 RT is even slower before longer/more complex episodes (e.g. those expected to have more rule switches; Schneider and Logan, 2006; Farooqui and Manly, 2017).

The subsuming nature of these programs is evidenced by the absence of switch cost across task episode boundaries. When task episodes consist of two or more types of task items (e.g. responding to shape and color of stimuli) such that any executed item can repeat or switch from the previous one, the expected switch cost reflected in higher RT/error rates for switch trials (Monsell, 2003) is absent for consecutive items executed as parts of different task episodes (Schneider and Logan, 2015). The change in the program at episode boundaries refreshes the lower-level item related cognitive configurations nested within it, causing no advantage of repeating a task item across episode boundaries. Subsuming nature of programs is also evidenced by the higher and more widespread activity elicited at the completion of task episodes compared to subtask episodes (Farooqui et al., 2012). Completion of subtasks only dismantles programs related to it, leaving the overarching task-related programs intact. In contrast, task completion dismantles programs at all levels.

Because attention (and control) always take place in the context of task episodes it is possible that deactivations previously attributed to them is actually related to the load of programs subsuming the task episode in which attention takes place. To test this we looked at activity across task episodes made of sequential events that were identical in their attentional demands. Deactivation is related to attention should be identical across these events. In contrast, deactivation related to the load of the subsuming program should be maximal at the beginning of the episode because the program at this point contains elements related to the entire length of the ensuing task episode and hence its load is maximal. This deactivation should then decrease as parts of the episode get executed, and the related elements within the program dismantle, decreasing the program load, e.g. at step 1 of a task episode made of four steps the program will contain elements related to all four, but at steps 2, 3 and 4 it will only contain elements related to the remaining three, two and one step, because those related to the earlier steps will have dismantled causing the program load to decrease gradually across sequential parts of the episode. Furthermore, if deactivation is not caused by attention to external stimuli, it need not be limited to regions that process internally generated information (i.e. the DMN).

Across three different kinds of task episodes we show that deactivations were indeed maximal at the beginning and decreased gradually across the episode. Further, this pattern of deactivation was not limited to the DMN but was shown by all regions uninvolved in processing the task content of the episode.

## Methods

### Experiment 1

Participants performed a rule-switching task (Figure 1). On each trial a letter and a digit were simultaneously presented for 1 s inside a colored margin. The color of the margin around the stimuli instructed which pre-learned rule to apply: blue - categorize the letter as vowel/consonant; green – categorize the digit as even/odd. In both cases they pressed one of the two buttons using index/middle finger of the right hand. Rule was determined at random for each trial.

**Figure 1.**
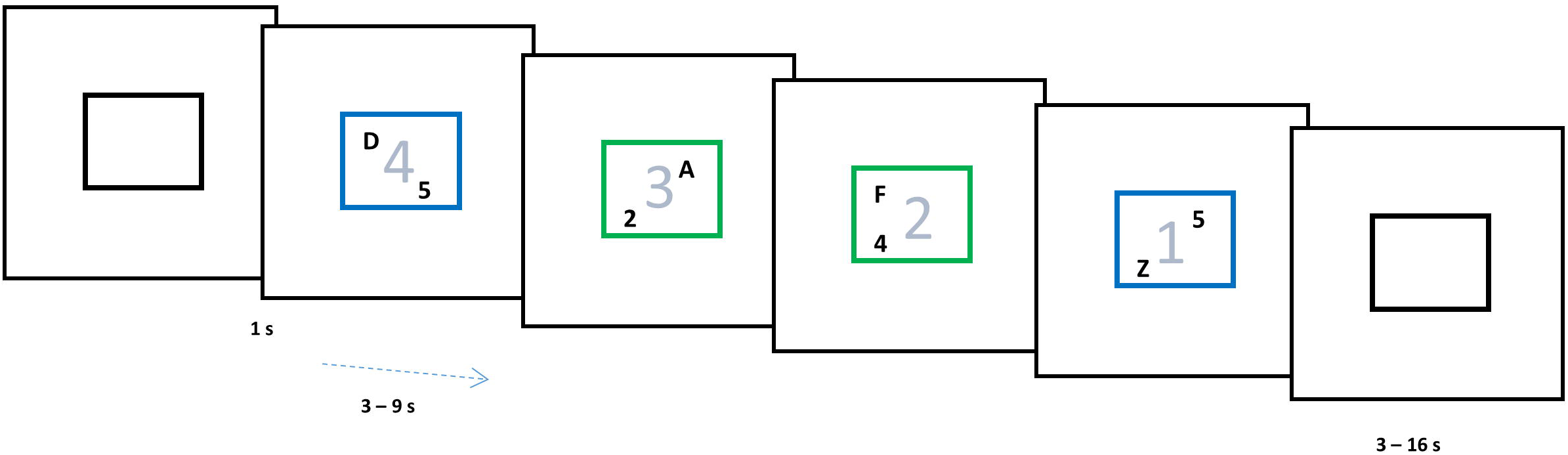
The color of stimulus margin determined if the letter was to be categorized as vowel/consonant (blue) or the number as even/odd (green). Subjects were biased to construe a set of four consecutive trials as one task episode. Trials of an episode had an irrelevant number in stimulus background that changed 4-3-2-1 across them and were preceded and succeeded by black margins.

Crucially, participants were biased toward construing each set of four consecutive trials as one task-episode by a recurring, trial-synchronized 4-3-2-1 background digit countdown that was otherwise irrelevant to the ongoing switching-task. The end of each task-episode was further signaled by the margin turning black. Inter trial interval varied between 3 to 9 (average 5.5) s. Participants did a total of 100 trials or 25 trial episodes.

All stimuli were presented at the center of the screen, visible from the participant’s position in the scanner via a mirror mounted within the head coil. The outer square margins surrounding the letter and the number subtended a visual angle of around 2°. The letter and number were presented in Arial font (size 30) and could appear at any of the corners of the outer square. Other than the 4- trial episodes described here, the experiment also involved 8-trial episodes. However, their analysis presents a further detail of the phenomenon described here which we deal with in another upcoming paper.

Note that individual trials were separated by long intervals (up to 9 s) and were independent of each other, and could be executed perfectly well without being construed as parts of a larger task episode. Participants were merely biased into construing a run of four successive trials as one task episode. Hence it is possible that participants may at times execute trials not as parts of an overarching task episode but as independent standalone trials. We reasoned that if a phenomenon is true for such weak and merely construed task episodes, it is likely to be also true for extended tasks that can only be executed as one task entity. We show that such is the case through experiments 2 and 3, where we also test additional predictions.

Eighteen (11 females; mean age, 25 ± 4.1years) right handed participants `with` normal or corrected vision were recruited. Informed consent was taken, and the participants were reimbursed for their time. The study had the approval of Research Ethics Committee. fMRI data were acquired using a Siemens 3T Tim Trio scanner with a 12 channel head coil. A sequential descending T2*- weighted echo planar imaging (EPI) acquisition sequence was used with the following parameters: acquisition time, 2000 ms; echo time, 30 ms; 32 oblique slices with slice thickness of 3 mm and a 0.75 mm interslice gap; in-plane resolution, 3.0 × 3.0 mm; matrix, 64 × 64; field of view, 192 mm; flip angle, 78°. T1-weighted MP RAGE structural images were also acquired for all participants (slice thickness, 1.0 mm; resolution, 1.0 × 1.0 × 1.5 mm; field of view, 256 mm; 160 slices). Experimental tasks started after 10 dummy scans had been acquired. These were discarded from the general linear model to allow for T1 equilibration effects.

#### Analysis

The fMRI data were analyzed using SPM8 (experiments 1 and 2) and SPM12 (experiments 3) using the automatic analysis pipeline (Cusack et al., 2015). Before statistical analysis, all EPI volumes were slice-time corrected using the first slice as a reference and then realigned into a standard orientation using the first volume as a reference. These realigned volumes were then normalized into the Montreal Neurological Institute (MNI) space and spatially smoothed using an 8 mm full-width half-maximum Gaussian kernel. During the normalization stage, voxels were resampled to a size of 3 × 3 × 3 mm. The time course of each voxel was high pass filtered with a cutoff period of 90 s.

Statistical analysis was carried out using general linear models (GLMs). We analyzed the data using two kinds of general linear models (GLMs). In the first, trials 1 to 4 of the construed episode were modeled with separate event regressors of 1 s duration. These were convolved with a basis function representing the canonical hemodynamic response (HRF), and entered into a general linear model with movement parameters as covariates of no interest. Parameter estimates for each regressor were calculated from the least-squares fit of the model to the data. We looked at regions where a linear contrast along the four trials (weighted [-3 -1 1 3]) for increasing activity across them. Contrasts from individual participants were entered into a random effects group analysis. In the second GLM we modeled 32 seconds of activity following the beginning of the episode with 16 two-second long finite impulse regressors (FIRs; Glover, 1999). This allowed us to derive an estimate of the time-course of activity across the duration of the construed task episode.

Other than events of interest, movement parameters and block means were included as covariates of no interest. All whole-brain results are reported at a threshold of p < 0.05 and corrected for multiple comparisons using the false discovery rate. Coordinates for peak activation are reported using an MNI template.

Regions of interest (ROIs) were created as spheres of 10 mm radius. These (in MNI space) were bilateral inferior frontal sulcus (IFS; central coordinate ±41 23 29), bilateral intraparietal sulcus (IPS; ± 37 56 41), bilateral anterior insula extending into frontal operculum (AI/FO; ± 35 18 3), anterior cingulate cortex (ACC; 0 31 24), and presupplementary motor area (pre-SMA; 0 18 50), all taken from Duncan, 2006; bilateral anterior prefrontal cortex ROIs (APFC; 27 50 23 and ±28 51 15) were taken from Dosenbach et al., 2006; posterior cingulate cortex (PCC; -8 -56 26) and Anterior Medial Prefrontal cortex (aMPFC; -6 52 -2) were taken from Andrews-Hanna et al., 2010. Coordinates for hand primary somatosensory area used in Experiment 1 were taken from Pleger et al., 2008. Primary auditory cortex ROI was made using probabilistic maps in the anatomy toolbox of SPM 8. ROIs were constructed using the MarsBaR toolbox for SPM (http://marsbar.sourceforge.net). Estimated data were averaged across voxels within each ROI using the MarsBaR toolbox, and the mean values were exported for analysis using SPSS.

## Results

As expected of task episodes, trial 1 had the longest RT (F_3,51_ = 11, p <0.001; inset Figure 2). RT on subsequent 3 trials did not differ (F_2,34_ = 0.8, p =0.4; linear contrast: F_1,17_ = 0.004, p = 0.9), showing no change in performance across them. Error rates across the four trials (5.7 ± 1.2, 3.0 ± 1.6, 4.8 ± 1.8, 5.1 ± 1.9) did not differ either (F_3,51_ = 1.6, p =0.2; linear contrast F_1,17_ = 0.004, p= 0.9). For a more detailed analysis of behavior across such task episodes see xxxxx, 2018^1^. Whole brain render in Figure 2 shows the set of regions where activity increased across the four trials of the construed episode. This was the case in regions identified with the Default Mode network (DMN) - medial prefrontal cortex (MPFC), posterior cingulate (PCC), cuneus, temporoparietal junction (TPJ) – along with right somatosensory and motor cortices; right superior and inferior frontal gyrus. In these regions trial 1 was accompanied by a deactivation and trial 4 with maximal activity, with trials 2 and 3 having intermediate levels of activity.

**Figure 2.**
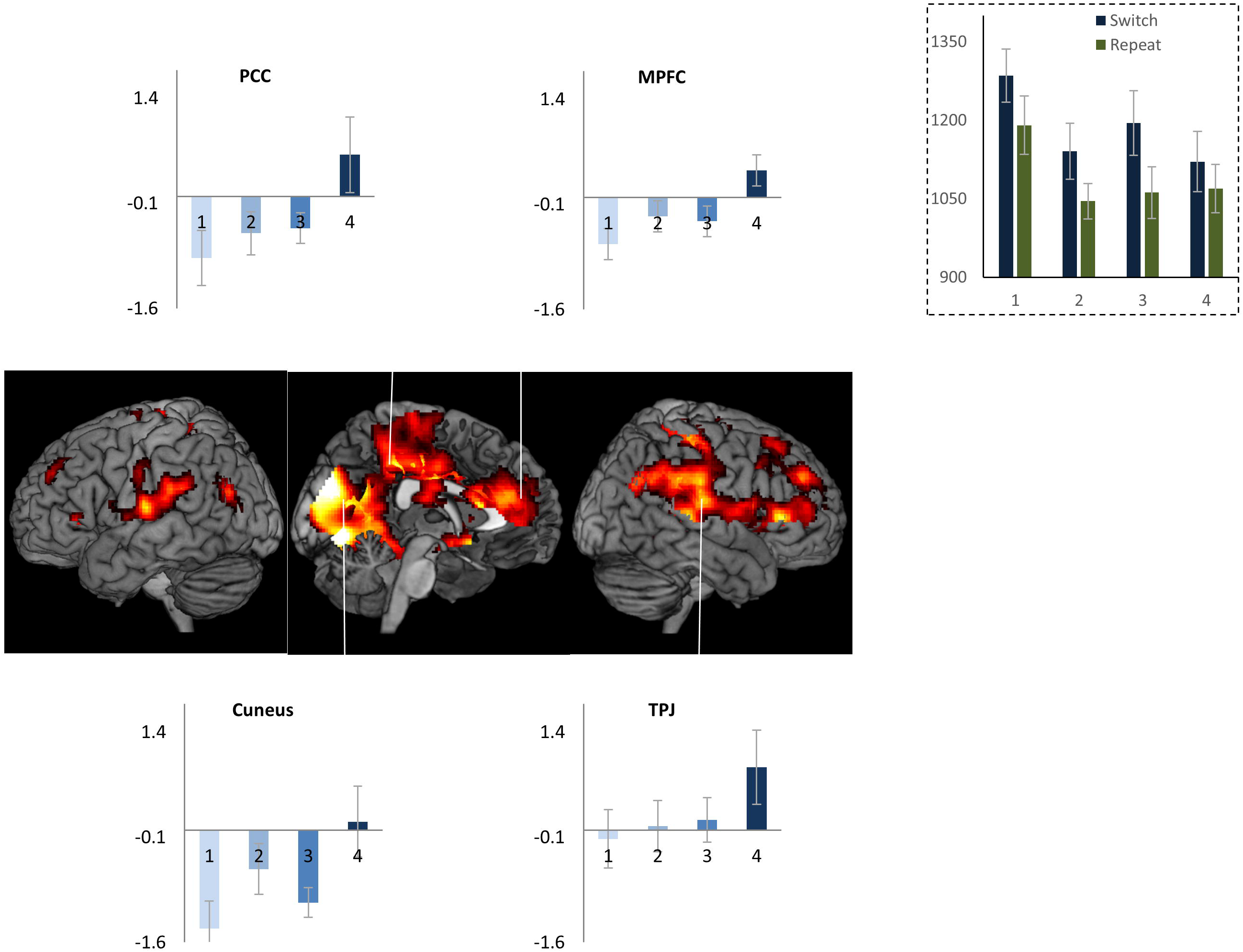
Inset shows reaction times across the four trials of an episode. Note that trial 1 RT is the highest; RTs across trials 2 to 4 do not differ. Whole brain render shows regions where activity changed across the four trials - medial occipital regions, cuneus, precuneus, posterior cingulate (PCC), retrosplenial, parahippocampal cortices, temporoparietal junction (TPJ), inferior parietal lobe, right somatosensory and motor regions, medial prefrontal cortex (MPFC), anterior cingulate, right inferior and superior frontal gyri extending onto right anterior prefrontal regions. Linked bar plots show the actual pattern of activity across these trials in some of the Default Mode regions. Error bars here and in all subsequent figures represent 95% confidence intervals.

We then conducted a region of interest (ROI) analysis of Multiple Demand (MD) regions - brain areas that are widely observed to be active (“task-positive”) across a range of challenging cognitive tasks (Fox et al., 2005; Duncan, 2006). While these are commonly thought to be anticorrelated to the DMN, we found that some of them – ACC and right APFC (main effect of trial position: F_3,51_>3.1, p <0.03; linear contrast: F_1,17_ > 3.8, p<0.06), with strong trends in Left APFC, bilateral insula and right IFS (p ≤ 0.1) – showed the same pattern of change in activity across the task episode as the DMN regions (Figure 3). In contrast, other MD regions – left IFS, bilateral IPS and pre-SMA – showed positive or non-differential activity across the four trials.

**Figure 3.**
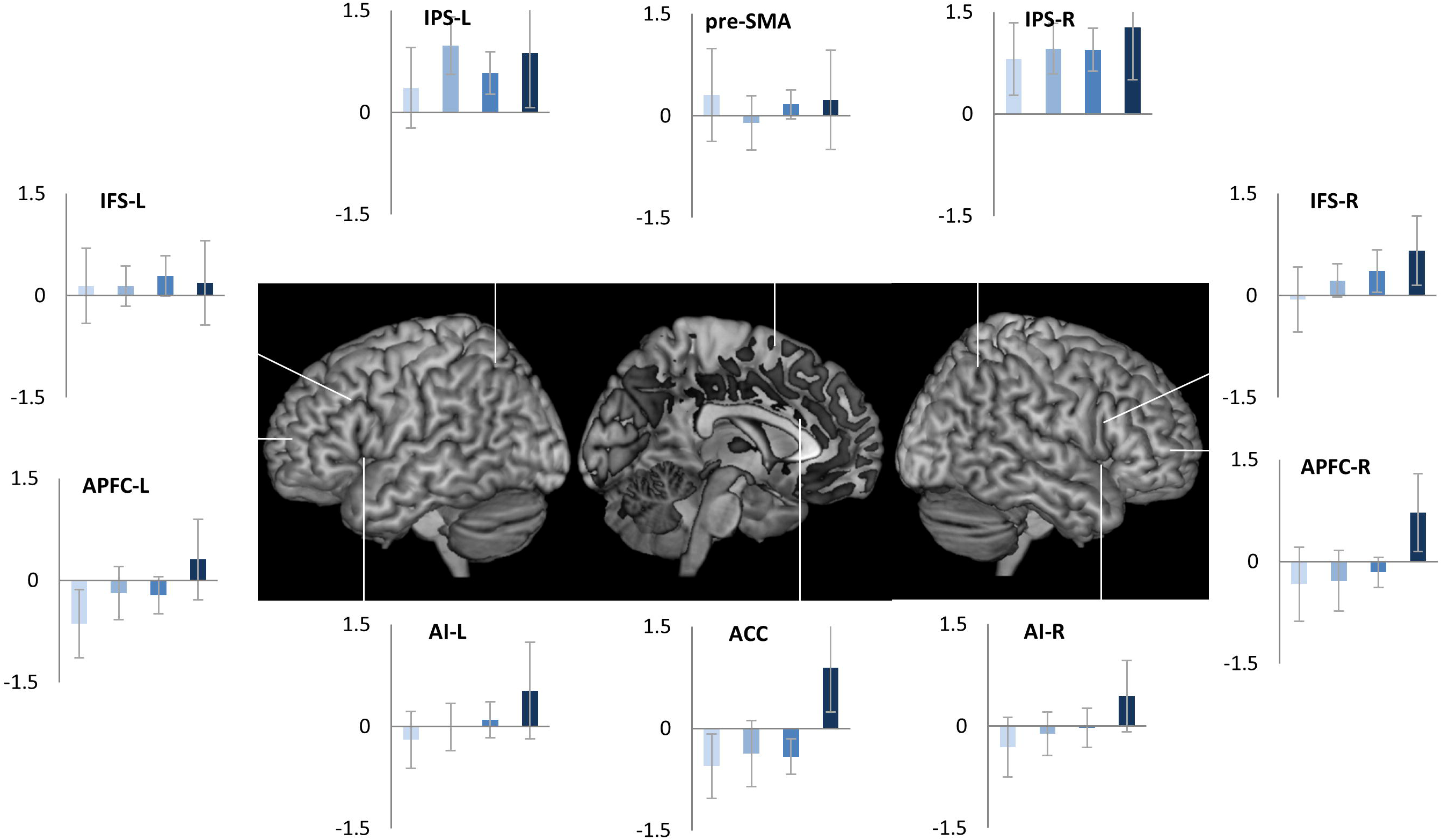
Of the cognitive control related fronto-parietal regions bilateral APFC and ACC showed differential activity across the four trials of the episode similar to that seen in DMN regions (figure 2). In contrast, IPS, pre-SMA and left IFS showed non-differential activity across the four trials.

These patterns were also evident in the estimates of time-course of activity across the task episode (Figure 4). DMN ROIs like PCC, aMPFC, and MD ROIs like APFC and ACC showed an initial deactivation starting at the beginning of the episode (i.e. trial 1 onset) followed by gradual return to the baseline as other trials were executed, with the activity reaching the baseline (captured by the estimates of the 1^st^ FIR regressor) by around 26 seconds (the average total duration of the episode was 24 seconds). In contrast, other MD ROIs (left IFS, IPS and pre-SMA) showed a very different pattern of activity with no deactivation during the episode.

**Figure 4.**
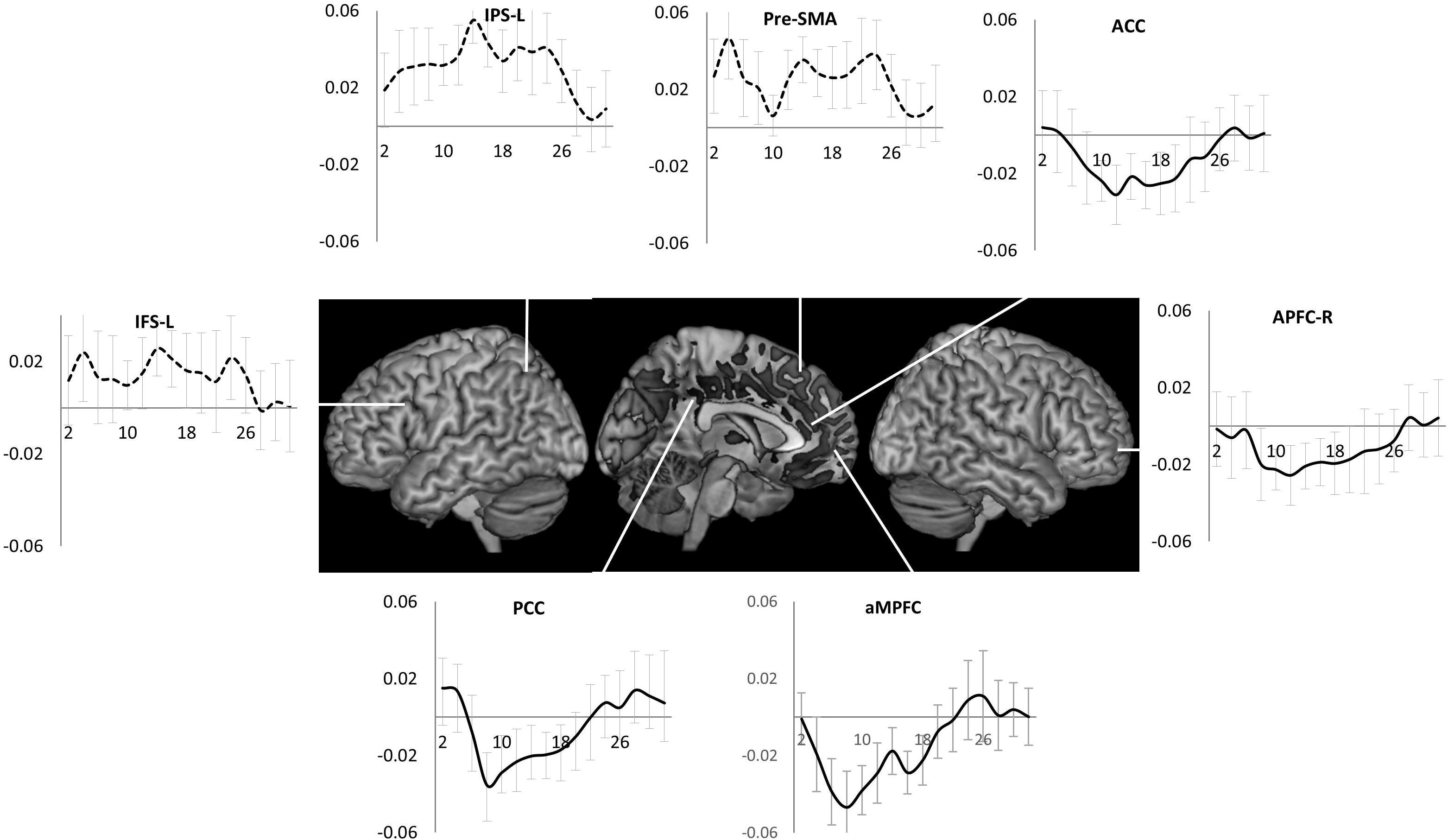
Time course of activity corresponding to the construed task episode in various ROIS. In regions where the four trials elicited differential activity – PCC, aMPFC, ACC and APFC-R - beginning of the episode caused a deactivation that returned to the baseline across the episode. This is in contrast to other fronto-parietal regions that did not show this pattern of deactivation followed by return to baseline (dotted line graphs), x-axis is time (s), error bars represent 95% confidence intervals.

The observation that the above pattern of deactivation was shown by DMN as well as the right motor regions (that will control the left side of the body) suggested that it may be present in regions *not* involved in executing individual trials making up the task episode. To investigate this thesis further, we identified regions controlling trial rule switches. These would show higher activity during switch trials (i.e. trials where the rule changed from letter to digit or vice versa) compared to repeats. These, shown as hot-spots in Figure 5, included left motor, left IFS, right middle frontal gyrus, pre-SMA extending into ACC, and left IPS. None of them showed the above pattern of deactivation across the task episode (bar graphs in Figure 5). We then identified regions that showed deactivation to task-switches. These (cold-spots Figure 5) were in medial prefrontal regions, posterior cingulate, precuneus, bilateral temporo-parietal junction, right IFS, right premotor and motor regions. Of these we arbitrarily selected four clusters adjacent to the hot-spots. All of them showed the typical initial deactivation followed by gradual activation across the episode (main effect of trial position: F_3, 17_ > 3.8, p < 0.02; linear contrast across trials: F_1,17_ > 4.04, p < 0.06). The difference across these two groups of ROIs is also evident in their time-courses of activity (line plots in Figure 5).

**Figure 5.**
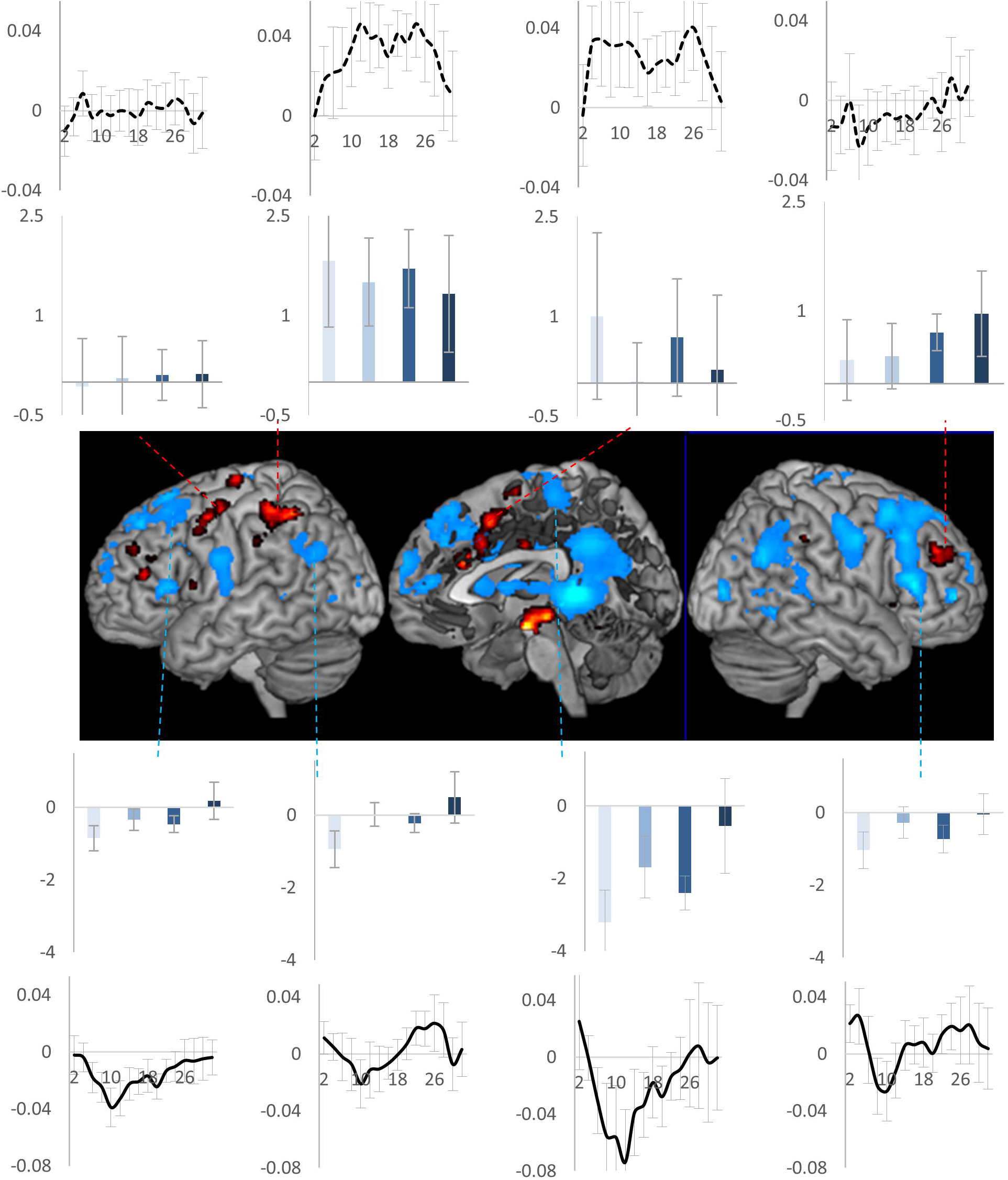
Activity across task episodes in switch trial positive (hot) and switch-trial negative (cold) clusters. Only switch-trial negative clusters showed the particular pattern of deactivation across the task episode. Bar plots show activity across the four trials, line plots show the time course of activity across the task episode. Error bars represent 95% confidence intervals.

We next examined the pattern of activity across task episode in two other regions expected to be unengaged in executing individual trials - right primary somatosensory hand region and bilateral primary auditory cortices (trials did not involve any listening nor any left hand action). These were contrasted with the left primary somatosensory hand region (expected to be involved in making right had button press). As shown in Figure 6, the uninvolved regions - bilateral primary auditory and right somatosensory hand region showed the sequentially changing deactivation (main effect of trial position: F_3,51_ > 5.0, p < 0.02; linear contrast across trials: F_1,17_ > 4.9, p <0.04) while the engaged left somatosensory hand region did not (L vs. R somatosensory difference F_3,51_ = 9.8, p <0.001). Again, the uninvolved regions showed initial deactivation followed by return to the baseline across the construed task episode.

**Figure 6.**
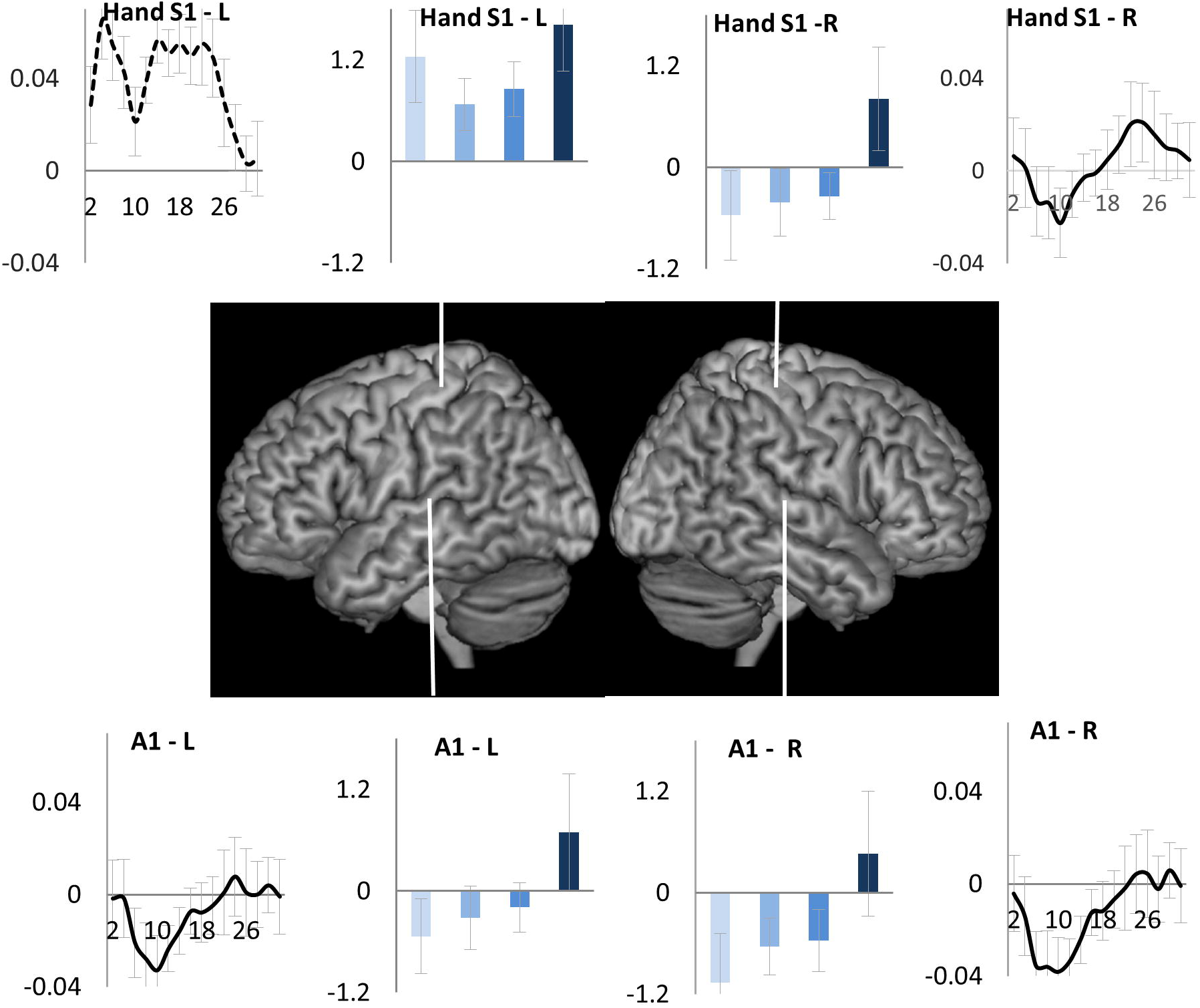
Regions uninvolved in executing individual trials - right somatosensory and bilateral primary auditory cortices (Al) showed deactivation elicited by the beginning of the episode that decreased and returned to baseline across the duration of the episode. In contrast, the engaged left somatosensory hand region showed a sustained activation across the episode.

Any account of deactivation that disregards the presence of task episode related programs would have predicted constant deactivation across all trials because participants essentially executed a flat sequence of identical trials with identical attention and control demands. In contrast, deactivation was *elicited* at trials that were construed as beginning a task episode. This deactivation then decreased as the construed episode was executed and reached baseline around the time of completion of the episode (Figure 4), with the last trial eliciting a positive activity. This pattern of deactivation can only be explained in relation to some task episode related cognitive entity – the program – that came into being at the beginning of the construed episode (eliciting maximal deactivation) and then decreased in its load across the episode (causing a gradual decrease in deactivation) and was dismantled at completion. This program related effect was present in the DMN regions previously seen to deactivate in response to cognitive load but was not limited to them, and additionally involved primary auditory, right somatosensory and motor cortices - regions expected to be uninvolved in executing trials making up the episode.

It may be pointed out that 95% confidence intervals of trial 1 activity estimates overlap with those of trials 2 and 3, and only trial 4 activity is significantly different from other trials. This may suggest that the deactivation elicited at trial 1 does not change across trials 1 to 3, with only trial 4 eliciting end-of-episode activity. While this is ruled out by the next experiment where activity does significantly increase between the first and the penultimate step, note that even this pattern can only be accounted through some episode related program. This is because trials here were standalone and their execution was not contingent on any episode related process like sustained attention or working memory that had to be maintained throughout the episode. That four of them constituted an episode was only in participants’ construal. There was no continuous performance, sustained attention, working memory or other process that participants started at trial 1 and ended at trial 4. Hence, whatever caused deactivation from trial 1 to trial 4 had to follow from the construal of these trials as one task episode. It’s also worth noting in this regard that the FIR estimates of time-series show that significant return towards baseline compared to the early nadir do occur before the end of the episode. Activity estimates from PCC and aMPFC (Figure 4), regions deactivating during rule switches (Figure 5) and primary auditory and right primary somatosensory hand region (Figure 6) show that activity has significantly risen above the nadir before the 24^th^ second (time of episode completion).

To see the generalizability of results of this experiment, we next investigated a task episode with very different structure and content. The task episode now involved covertly monitoring sequential letter presentations. Since this would be less demanding than executing the rule switch trials of experiment 1, more regions would be uninvolved in controlling/executing individual steps, and hence more widespread regions would show gradually decreasing deactivation across the length of the episode.

### Experiment 2a

At the beginning of a trial episode participants (total 21, 15 females; mean age, 24.5 ± 4.1years) saw a 3 letter cue (e.g. DAT). They were to then monitor a sequence of 40 single letter presentations (1 s) for the occurrence of these targets (in the correct order). If the cue was ‘DAT’, they were to search for D, then A and then T. Searching for A was only relevant once D had been detected and searching for T was only relevant once D and A had been detected. At the end of each sequence participants were probed to report whether or not all three targets had appeared by pressing one of two buttons (right index/middle fingers). A complete set of all three target letters appeared in 50% of trial episodes. In the rest none, one or two targets appeared.

Detecting all three targets and answering the probe correctly increased the participants’ score by 1, otherwise the score remained the same. Note that for this experiment the phrases ‘trial episode’ and ‘task episode’ are synonymous because each trial was temporally extended and formed a task episode. The episode consisted of monitoring 40 letter events over a period of 40 s, then answering the probe at the end of that period. Across different instances, the 40 s long trial episodes could be organized into up to four phases – the three searches plus the passive wait between the third target detection and the probe (Figure 7). While the total length of the trial episode was fixed at 40 s, the length of any of the four component phases varied between 1 s to 40 s. Note that these phases should not be thought of as steps that had to be obligatorily executed to complete the task episode because the episode got completed irrespective of the number of phases completed. When, for example, only one target appeared in the trial episode, the episode got completed during phase 2. At the same time when averaged across the entire session the four phases corresponded to sequentially later parts of the trial episode.

**Figure 7.**
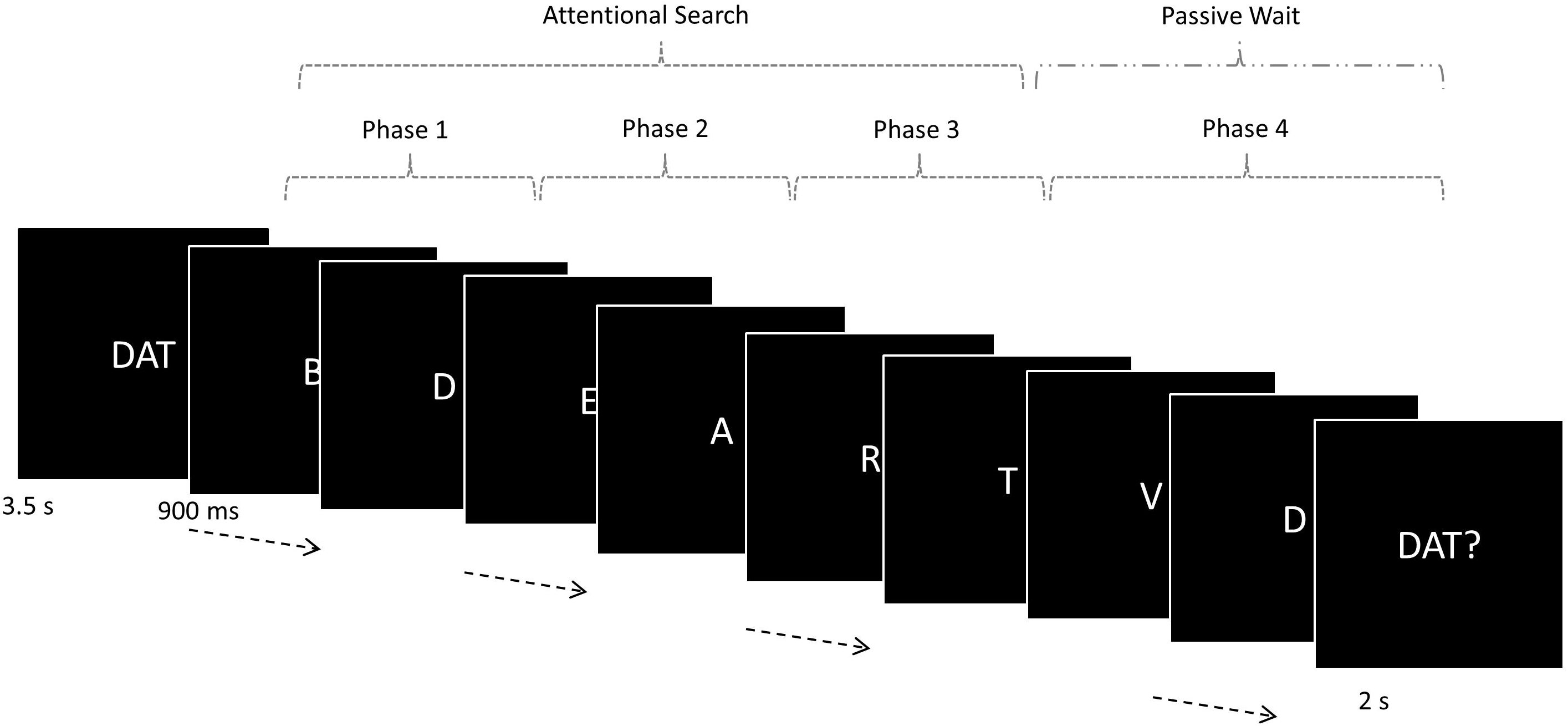
Trial episodes began with a pronounceable three-letter cue. The three letters of this cue were to be covertly detected, in the correct order, in the ensuing episode that involved covertly monitoring 40 letter sequential letter presentations. After all three had been detected, search stopped and subjects waited for the letter presentations to end, at which point a probe appeared asking if all three targets had appeared. The trial episode thus consisted four sequential phases – the first three of which involved visual search for a specific letter while the fourth involved a wait for the probe to appear.

All stimuli were presented at the center of the screen, visible from the participant’s position in the scanner via a mirror mounted within the head coil. Letters subtended a visual angle of 2° vertically. The experiment was controlled by a program written in Visual Basic. Participants learnt the task in a 10 min pre-scan practice session. The scanning session lasted an hour and was divided into three separate runs, each consisting of 14 trials.

The three target events (data not presented in current paper) were modeled with event regressors of no duration while the four phases along with the cue and probe were modeled using epoch regressors of width equal to their durations. All trial episodes were thus modeled. As in experiment 1 we did a linear contrast that looked for increase in activity across phases 1 to 4. To get the time-course of activity across the task episode, in a second analysis, we modeled 52 s of activity following the beginning of the episode with 26 FIRs. Cue, probe and target events were additionally modeled as epoch regressors of width equal to their durations and convolved with HRF. Other aspects of fMRI acquisition, pre-processing and analysis were identical to experiment 1.

## Results

Average response time was 727 ± 35 ms, while average accuracy was 95.2% (±1.1).We first identified areas involved in visual search. Activity in such areas would be higher during active visual search (phases 1 to 3) than during the period of passive waiting (phase 4). This was the case in a bilateral cluster that included parts of occipital pole and extended into the inferior division of lateral occipital cortex (Hot regions in Figure 8). Decreasing the threshold to uncorrected p < 0.05 showed an additional cluster in left frontal eye field. Other frontal and parietal regions did not show any significant cluster. In the light of experiment 1, remaining cortical regions can be expected to show change in activity across the length of the episode.

**Figure 8.**
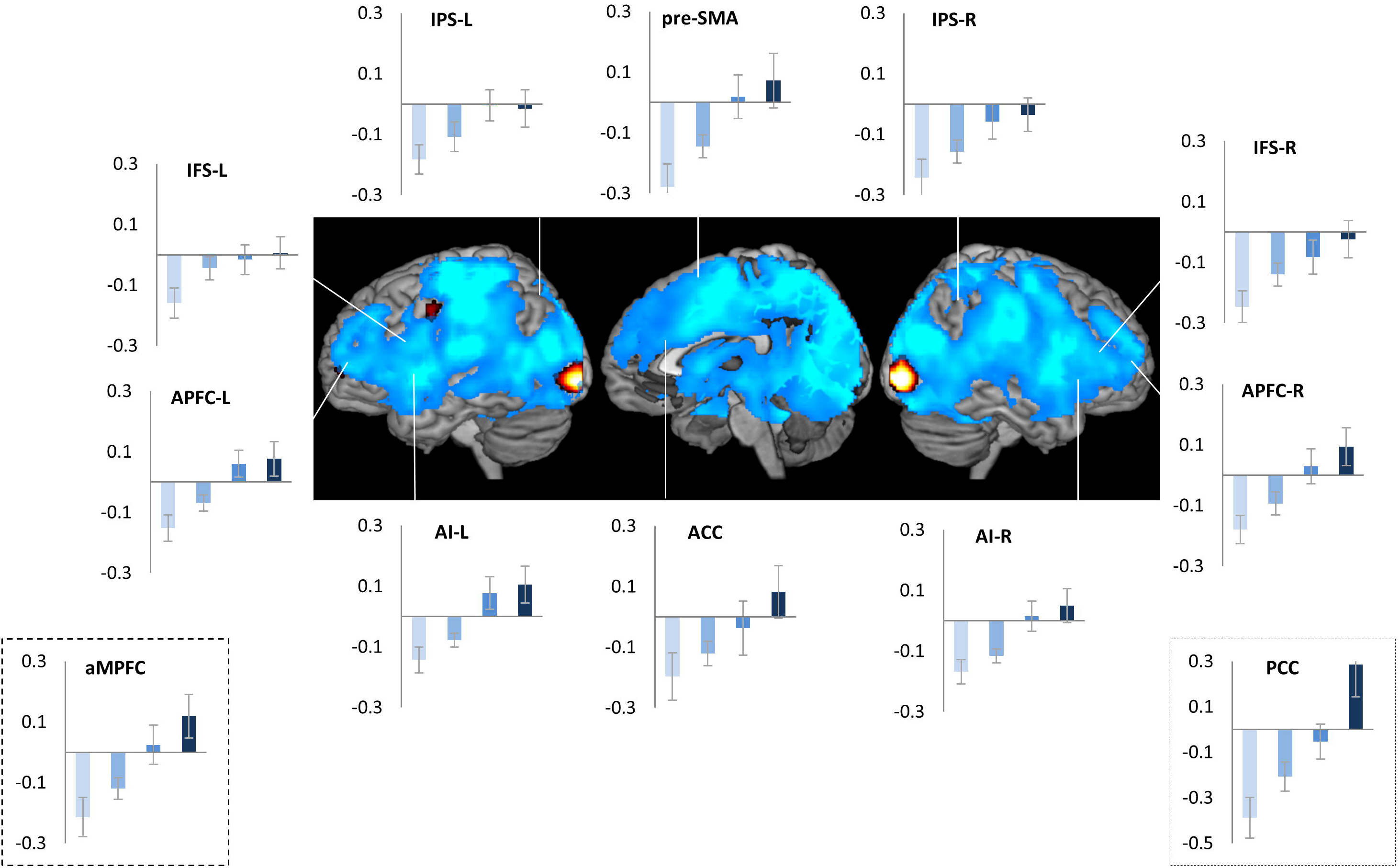
Hot render are the regions where activity increased during attentional search compared to passive rest. Cold render are the regions that showed sequential change in activity across the four phases of the episode. Note that the two renders do not overlap. Very widespread regions (cold render), with the notable exception of some visual regions, showed sequential change in activity across the four phases of the task episode. The pattern of activity (initial deactivation followed by stepwise activation) in all major MD regions was identical to that in the DMN regions (bordered plots). Error bars represent 95% confidence intervals.

Cold regions in Figure 8 show regions where activity increased across the sequential phases making up the trial episode. This pattern was present in very widespread regions that included all major DMN and MD regions as well as all non-visual sensory and motor cortices. Parts of cerebellum, basal ganglia, thalamus and medial temporal regions also showed an identical pattern. Thus, almost the entire brain with some islands of exception (including notably, middle occipital regions that were sensitive to visual search) showed increase in activity across the four phases of the trial episode.

The pattern of activity (plots in Figure 8, Figure 9) across the trial episode was similar to experiment 1 - initial deactivation followed by gradual return to the baseline across the duration of the episode, but this time reaching baseline (captured by the estimates of the 1^st^ FIR regressor) by the 40^th^ second (Figure 9), because the episode duration was 40 seconds. Additionally, as suggested by the whole brain results both MD and DMN regions showed the same pattern. Thus, despite the different structure and content of task episodes, experiment 2A showed the same pattern of gradually decreasing deactivation across the length of the task episode as in experiment 1. Additionally, this pattern was shown by both MD and DMN regions, as well as all sensory (except visual) and motor regions. This further supported the thesis that this pattern of deactivation during task episodes is shown not just by the DMN regions but by all regions uninvolved in executing component task items making up the episode.

**Figure 9.**
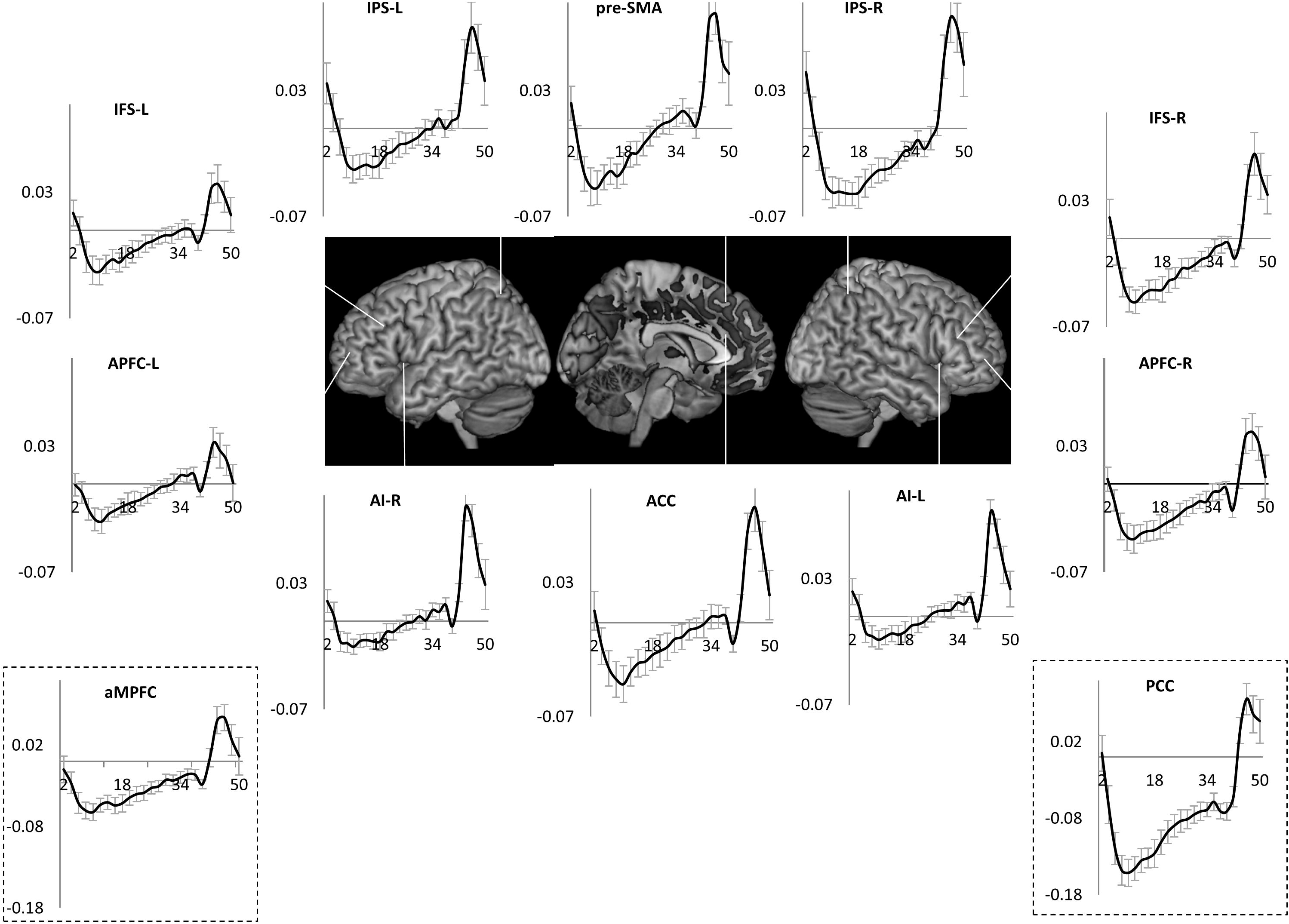
Time-course of activity across the task episode in various control related fronto-parietal regions. Error bars represent 95% confidence intervals. Note that the pattern was identical to that in a representative DMN regions (PCC). Across all of them, the beginning of the episode elicited a deactivation that gradually decreased across the length of the episode, with episode completion (at 40 s) eliciting a strong activation.

### Two kinds of Goals

Goals can be understood in at least two ways – (1) that which completes the task episode, (2) the intended aim whose achievement was sought through the task episode. While the two kinds of goals frequently coincide, they can be dissociated. One can go shopping with a list and complete the task episode of shopping (i.e. walk through all the aisles of the supermarket and check out) without finding everything on the list. Here goal 1 gets completed but not goal 2. The two were dissociable in the current experiment as well. All trial episodes consisted of seeing through 40 letter presentations, but in only half of them could all three targets be detected. Thus, while goal 1 (completion of the task episode) was achieved in all trial episodes, goal 2 (detecting all three targets) was achieved in only half of them. Further, one could move closer in relation to one goal type without moving much in relation to the other goal type. For example, in trial episodes where no targets appeared, goal 1 would complete before even the first step towards goal 2 had been completed. In contrast, when in some trial episodes all three targets appeared before the 10th letter event, goal 2 had completed while the participant had not even progressed halfway in relation to goal 1 (completing the trial episode).

Searches 1 to 3 for the three sequential targets represented sequential steps in relation to goal 2 and, when averaged across the experiment, also corresponded to sequentially later parts of the trial episode, hence the increase in activity across them in Figure 8 could be related to either of them. Deactivation could have decreased across them because participants were moving closer to the completion of the trial episode or because they were moving towards goal 2.

Programs are primarily related to the execution of task episodes, irrespective of whether a goal 2 gets achieved through that episode. Program instantiated at the beginning in the current experiment should therefore be related to the execution of the 40 s long trial episode, and the magnitude of the active program at any point in the execution of the episode should be related to the amount of trial episode that has been executed and the amount that remains to be executed. This predicts that the activity corresponding to the same search phase (e.g. search 2) should vary depending on its position within the trial episode. A search 2 that finishes before the 20^th^ letter event should be accompanied by greater deactivation than a search 2 that starts after the 20^th^ event. This is because the length of trial episode that remains to be executed is greater during the former than the latter, hence the magnitude of active program will be greater during the former than during the latter. Further, whether by the 20th letter event (the mid-point of the trial episode) the participant has completed up to search 2 or search 3, the magnitude of active program should roughly be the same because in both cases half of the episode remains to be executed. The program related activity should therefore be similar in the two cases.

To test these predictions we separately modeled instances of searches 2 and 3 that ended before the mid-point of the episode (early-search 2 and early-search 3) as well as instances of these searches that began after the episode mid-point (late-search 2 and late-search 3). Early-search 2 and early-search 3 began and ended at nearly identical positions within the trial episode. Same for late-search 2 and late-search 3. In contrast, early and late search 2 (as well as early and late search 3) corresponded to positions from the initial and later halves of the trial episodes. The magnitude of active program would be greater during early search 2 and 3 than during late search 2 and 3 because more of the episode remained to be executed during early search 2 and 3. The magnitude of deactivation should therefore be greater during early compared to late searches. At the same time the magnitude of deactivation may not differ between early searches 2 and 3 or between late searches 2 and 3.

We did a repeated-measures ANOVA comparing the effect of position in relation to goal 2 (search 2 vs search 3) and the effect of position within the trial episode (early vs late). As evident in Figure 10, deactivation during early searches was greater than during late searches in all MD and DMN ROIs examined (F_1, 20_ > 5.3, p <0.03), except APFC (L: F_1, 20_ = 1.9, p = 0.2; R: F_1, 20_ = 4.1, p = 0. 06). As expected, searches 2 and 3 did not significantly differ though it did show trends in some ROIs (F_1, 20_ < 3.9, p > 0.06). Interaction between these two effects did not reach significance in any ROI, though, again, it showed trends in IFS-R and APFC-R (F_1,20_ < 3.7, p > 0.07).

**Figure 10.**
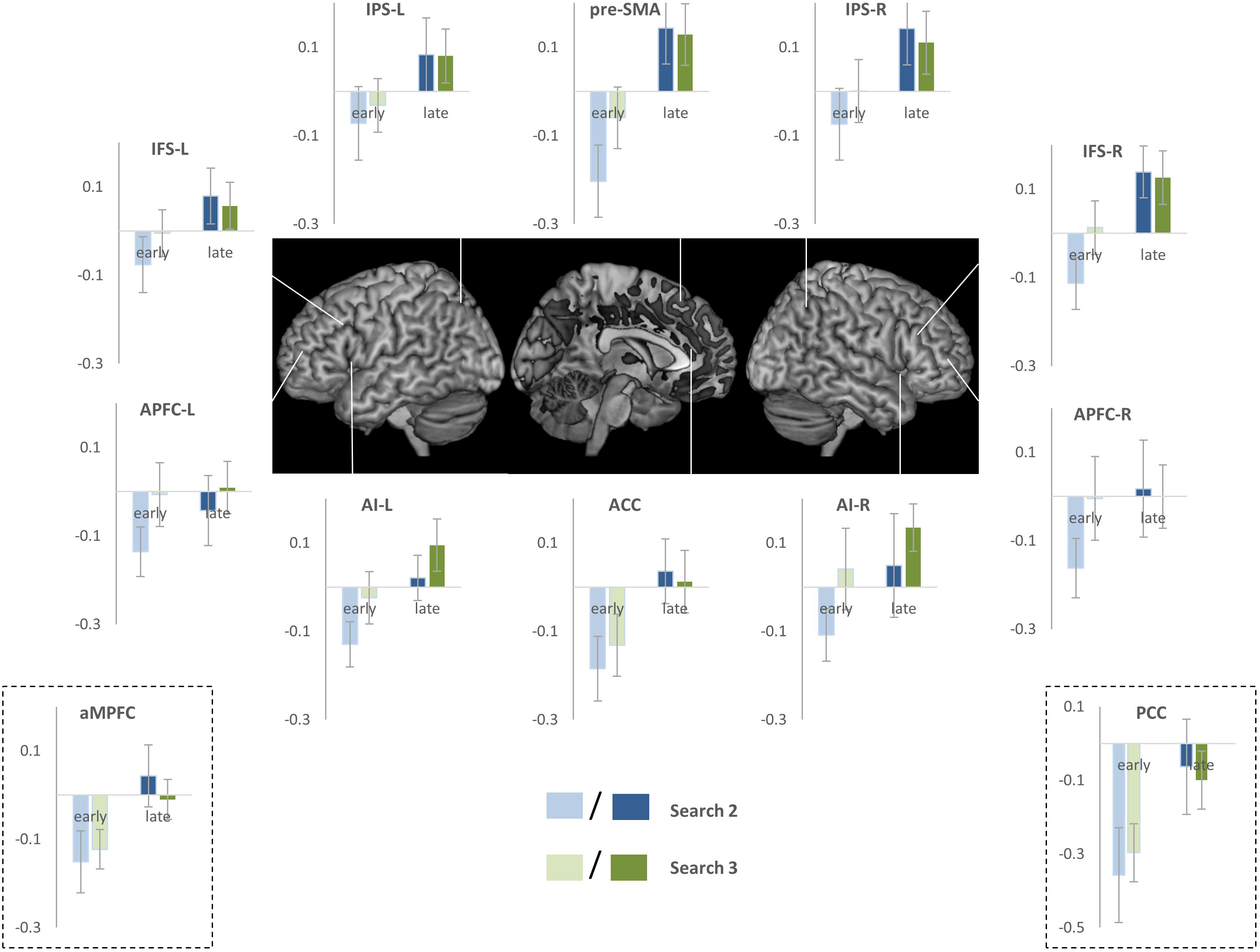
Estimates of activity during search phase 2 when it ended before the middle of the trial episode (early) compared with search phase 2 activity when it started after the middle of the episode (late). Note that activity in all ROIs during late search phase 2 was higher than that during early search phase 2.

#### Experiment 2B

It may be claimed that the initial deactivation is elicited by attentional/control demands and that the subsequent decrease in this deactivation across task episode is a result of weakening of attention/control due to continuous task execution. In this experiment we rule this out, and provide further evidence that the deactivation and its subsequent decrease result from episode related programs.

If the onset of attention caused initial deactivation then deactivation at the beginning of an 18 s long task episode should be equal to that at the beginning of an otherwise identical but 40 s long task episode. Furthermore, if weakening of attention due to continuous task execution caused a decrease in the magnitude of this initial deactivation, then the trajectory of activity time-course should be identical across these two task episodes for 18 seconds. This is because attentional decrement or any other purported effect of continuous task execution will be only be related to the time that has been spent on the task, and not to the length of the episode remaining to be executed. In contrast, if the deactivation and its subsequent decrease are related to the episode related program, time-course of activity will differ across the long and short episodes. First, because the deactivation results from the load of the program, which will be less for the shorter episode, the magnitude of initial deactivation will be less for the shorter episode. Second, because the decrease in this deactivation results from the dismantling of program elements related to completed parts of the episode, the return to baseline will be faster in shorter episode with activity reaching baseline around the expected time of completion.

Current experiment was identical to experiment 2A except that its trial episodes were shorter in length (8 to 18 s compared to 40 s). Seventeen participants (10 females; mean age 23.4 ± 6.4 years) executed 160 task episodes across 4 fMRI sessions. We modelled trial episodes that lasted 15 to 18 s with 12 FIR regressors, starting from the beginning of the trial episode and covering the subsequent 24 s period. Shorter episodes were separately modeled. Cue, probe and target events were additionally modeled as epoch regressors of width equal to their durations and convolved with HRF. Other aspects of fMRI acquisition, pre-processing and analysis were identical to experiment 2A.

## Results

Average RT was 750 ms (± 49); error rate was 3.7% (± 1.5). If deactivation and its subsequent decrease was caused by attention and its purported decrements it should be identical across the initial 18 s period of the trial episodes from the current and the previous experiment. In contrast, if the deactivation was related to the magnitude of the program then the initial deactivation should be stronger for the 40 s long trial episodes of the previous experiment, because the magnitude of program executing longer task episodes will be larger and have a greater cognitive load. Further, because the program in the current experiment dismantles over the 18 second period, its related deactivation should decrease and reach baseline at around 18 s. As is evident in Figure 11, this was indeed the case. The initial deactivation in this experiment was smaller in magnitude than in the previous one; activity in the current experiment returned to baseline around 18 s the expected time of completion. A repeated-measures ANOVA comparing time-courses of activity across the 18 s period of trial episodes from the current and the previous experiments showed significant difference across them in all ROIs (F_8,288_ > 3.9, p > 0.001) except ACC and left APFC, although it did show strong trend in left APFC (p =0.05).

**Figure 11.**
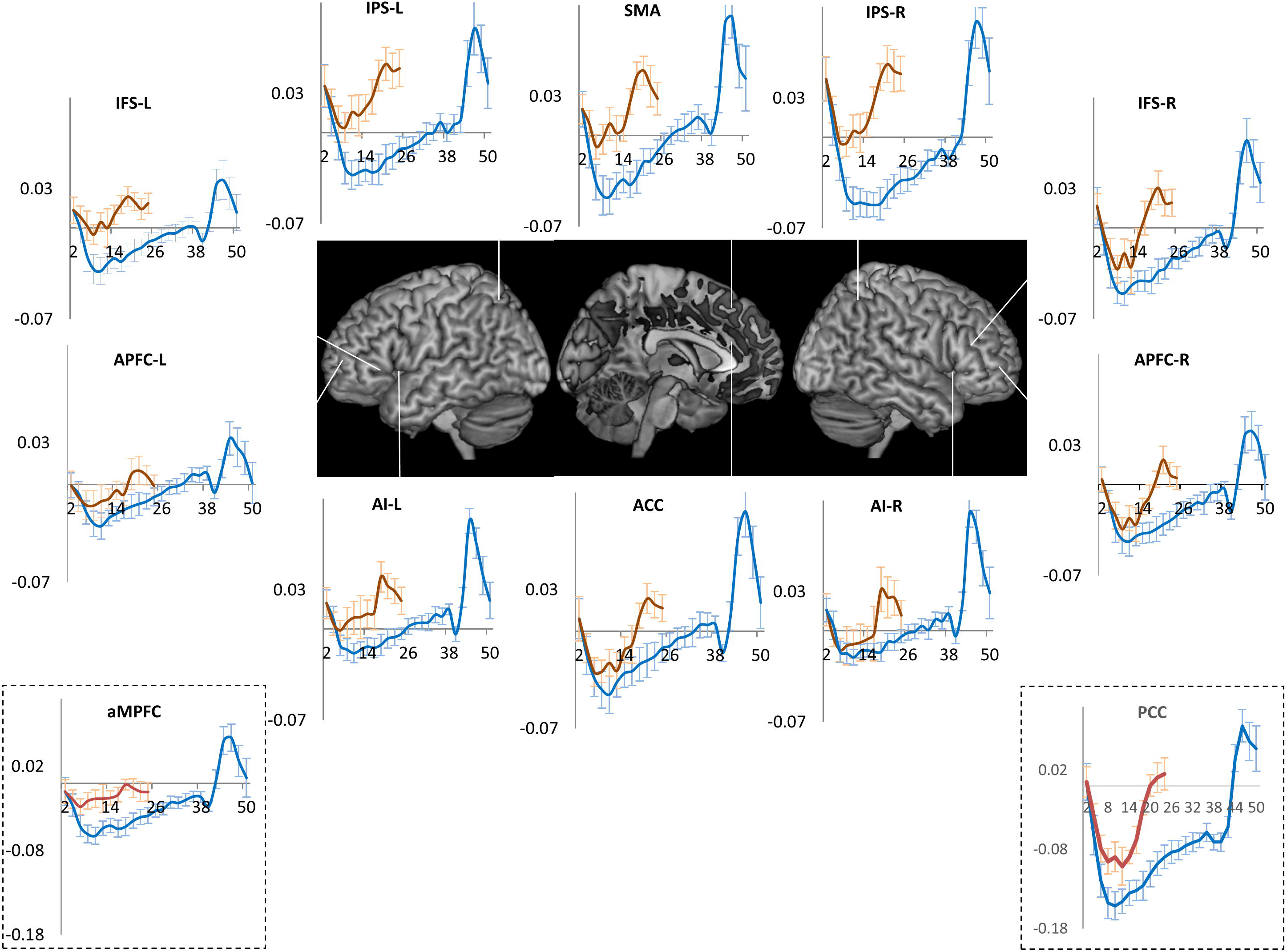
Orange line plots the time-course of activity across the 15-18 second long trial episodes of experiment 2B, blue lines plot the activity time-course across the 40 second long trial episodes of experiment 2A. Note in experiment 2B nadir is shallower and activity returns to baseline by the 18^th^ second, the expected time of completion.

### Experiment 3

In this experiment we decisively tested the thesis that deactivation during task episodes is shown not just by the DMN but potentially by all regions not engaged in performing the ongoing task. Seventeen participants (14 females; mean age 24.2 ± 5.3 years) executed two kinds of task episodes – Number and Shape (Figure 12), with right and left hands respectively. When they executed the Number episode with their right hand, the motor region controlling the idle left hand in the right hemisphere will be uninvolved, while the left hemisphere motor region controlling the right hand will be uninvolved when they executed the Shape episode with left hand. The gradually decreasing deactivation should therefore be shown by right but not left hemisphere hand region during the number episode and by left but not right hemisphere hand region during the shape episode.

**Figure 12.**
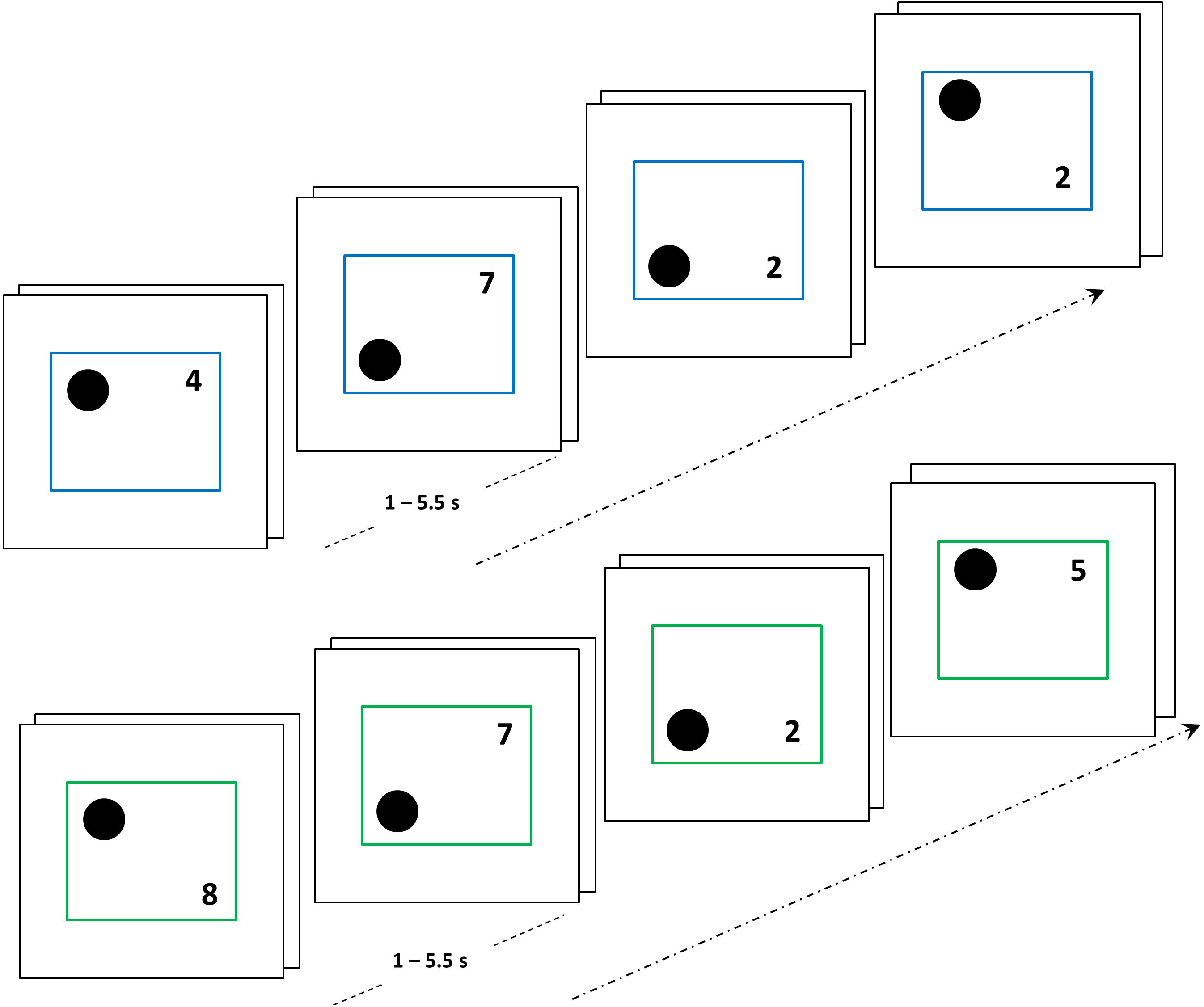
Episodes consisted of four steps, each of which involved a paired-trial i.e. two trials that followed without any interval. In a shape episode (blue margin trials) participants categorized the shape as above/below the center with their left hand, while in a number episode (green margin trials) they categorized the number as odd/even with their right hand.

The identity of the episode was cued by the color of stimulus margins (e.g. Number: Green, Shape: Blue). Participants executed a four step trial episode. Each step consisted of a pair of trials i. e. two trials juxtaposed without any interval. Stimuli consisted of a circle and a number on the left and right sides respectively of a surrounding square. These remained on screen till a response was made. On every trial of Number episodes participants indicated if the displayed number was even or odd using their right hand (index finger: even, middle finger: odd). During Shape episodes participants indicated if the black circle in the stimuli was above or below the center using their left hand (middle finger: up, index finger: down). Additionally, participants kept an internal count and after executing 4 phases (or 8 trials) pressed an extra button (the ‘end’ response) with the middle finger of their task relevant hand, i.e. left hand during shape episodes, right hand during number episodes. The internal count and the ‘end’ response were included to help ensure that participants construed the trials as part of a longer episode, not as a series of independent entities. To reinforce this, a positive feedback tone was presented if the ‘end’ response occurred within 500 ms. whilst a response > 500 ms. received a low pitched error tone.

The interval between consecutive steps was jittered from 1 to 7 s. The surrounding square remained on for the entire duration of the episode and went off at its completion. The interval between successive episodes was also jittered between 1 and 7 s. Participants executed a total of 30 task episodes consisting of equal number of shape and number episodes. The two episode types were randomly interspersed. After the main experiment session, participants executed a 10 minute localizer task. This consisted of 16 second long alternating blocks of face and place trials. On face trials they categorized face pictures as male or female; on place trials they categorized scenes as indoor or outdoor. Importantly, for one block type they used index and middle finger of right hand and for the other same fingers of left hand. The actual combination was balanced across participants. Through this we delineated somatosensory and motor voxels that were more active during right and left hand button presses. These can be expected to also be engaged in making respective button presses during the main experiment session.

Each paired-trial was modeled with an epoch regressor that extended from the stimulus onset for the first trial of the pair to the response of the second trial of the pair. We then did a linear contrast of the estimates of activity from the four paired-trials looking for regions where activity increased linearly across them. For the second GLM we modeled 34 seconds of activity following the onset of the task episode with 17 two-second long FIRs. In localizer session of experiment 3 the right and left hand executed blocks were modeled with epoch regressors equal to their length. Estimates of activity were contrasted to get regions where activity during the right hand executed blocks was higher than the left hand executed blocks and vice-versa. The two hand region related voxels of interest were delineated for every participant individually. Other aspects of fMRI acquisition, pre-processing and analysis were identical to experiment 1.

If the deactivation followed by increase in activity across the task episode is shown by regions not involved in executing component task items then during the number episodes this pattern will be shown by the left hand related voxels lying in the right hemisphere, and during shape episodes by the right hand related voxels lying in the left hemisphere. Regions like PCC and MPFC (DMN regions) and primary auditory cortices (part of the uninvolved sensory regions) will show this behavior during both task episodes.

## Results

As evident in Figure 13, first trial of episodes had the longest RT (F_7, 112_ = 103, p < 0.001). The first trial of each paired-trial also had longer RT than the second trial, because it began a sub-episode consisting of two trials (De Jong, 1995). Number episodes had higher RT than shape episodes (F_1, 16_ = 16, p =0.001), but the pattern of RT across sequential positions did not differ between them (F_7, 112_ = 0.7, p = 0.6). Error rates too were higher at trial 1 (F_7, 112_ = 3.2, p = 0.004). RTs across paired-trials 2 to 4 were marginally different (F_2, 32_ = 3.3, p = 0.05; linear contrast: F_1,16_ = 3.3, p =0.09) because the first trial of paired-trial 2 was slower than the first trials of subsequent paired-trials (0.91 ± 0.03 s, 0.85s ± 0.04 s, 0.88s ± 0.04 s on paired-trials 2, 3 and 4 respectively). Error rates did not differ across them (F_2, 32_ = 0.02, p = 0.8; linear contrast: F_1,16_ = 0.02, p = 0.9).

**Figure 13.**
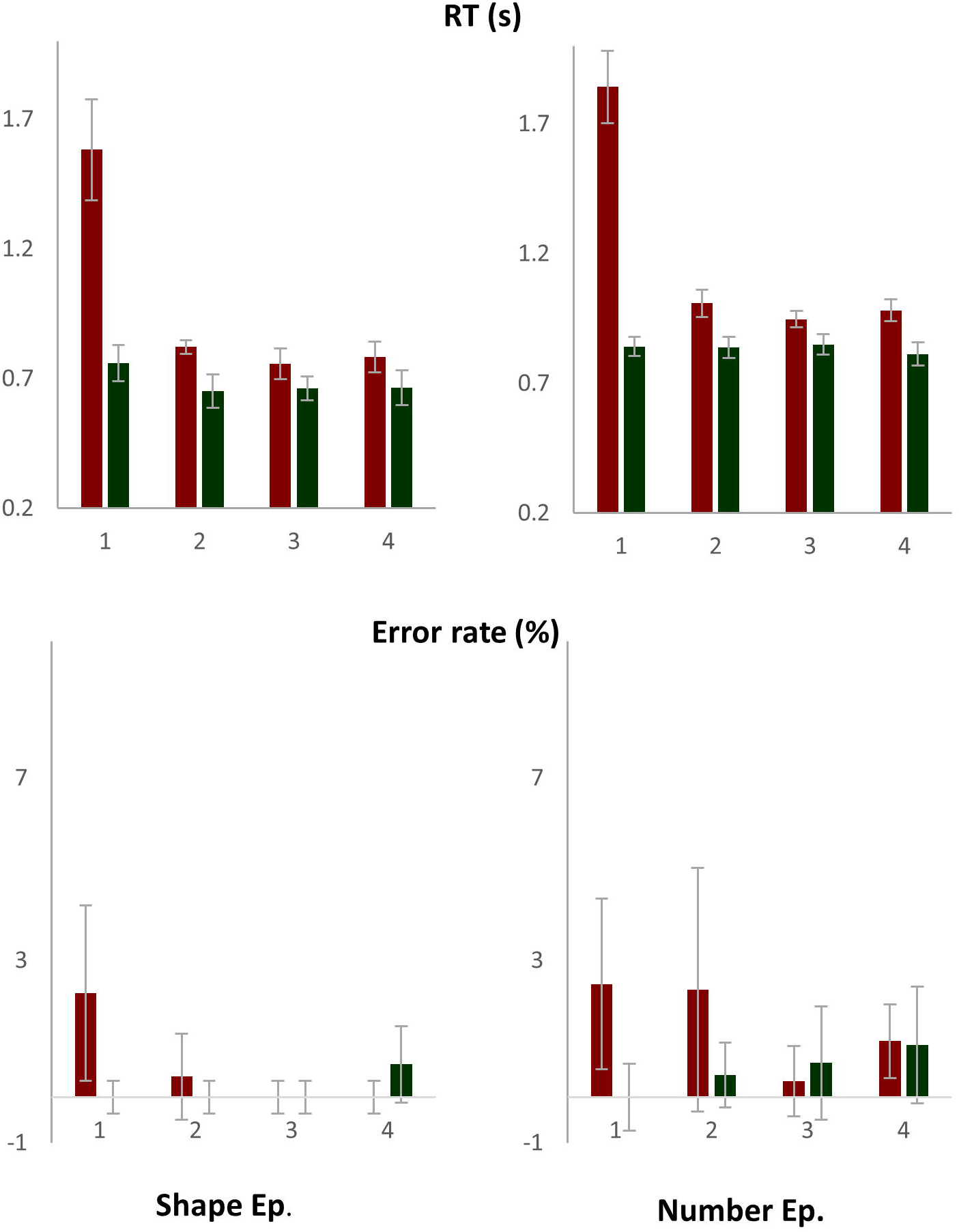
The first trial of the episode had the longest RT and higher error rates. The first trial of each paired trial also had higher RT than the second trial. RTs and error rates did not increase across steps 2 to 4.

As apparent from the graphs of Figure 14, the hand sensory-motor region in left hemisphere controlling the right hand showed initial decrease followed by gradual increase during the left hand executed Shape episodes. In contrast, the hand in right hemisphere controlling the left hand this pattern during the right hand executed Number episodes. Regions like PCC and aMPFC (DMN regions) and primary auditory cortex (an uninvolved sensory region) showed this pattern during both episodes. The whole brain render in Figure 14 shows regions where a linear contrast looking at increase in activity across the four phases was significant during Number (blue) and Shape (green) episodes. Regions in cyan were significant for both episodes. Note that most DMN regions along with right superior and inferior frontal gyri are cyan in color.

**Figure 14.**
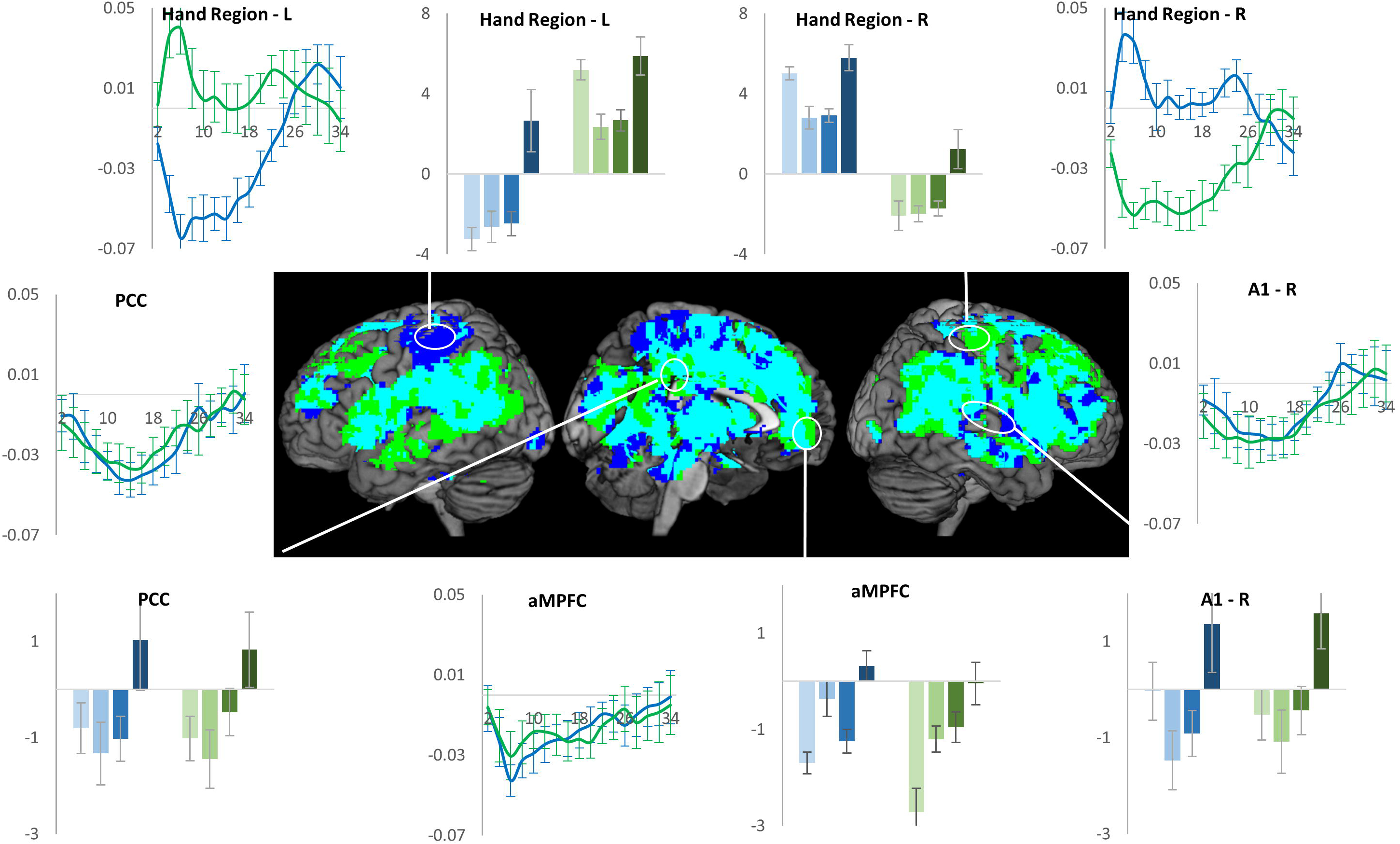
During the right hand executed Number episodes the uninvolved right hemispherical hand region showed initial deactivation followed by gradual increase (green plots). In contrast, during the left hand executed Shape episode this pattern was shown by the uninvolved left hemispherical hand region (blue plots). DMN regions – PCC and, and primary auditory cortex (a region expected to be uninvolved during both episodes) showed this pattern during both task episodes. Whole brain render shows regions where linear contrast of activity across the four trials was significant: green - number episodes, blue - shape episodes, cyan - both.

## General Discussion

Across three different experiments with task episodes differing in length, structure and content, widespread regions showed maximal deactivation at the beginning of task episode execution followed by gradual return to baseline as parts of the episode were executed, with the activity reaching baseline towards the end of the episode. This pattern of deactivation cannot be explained by changes in attention or other control demands related to task related perception, rule selection, decision making and response selection, because these remained constant across the duration of the episodes. Neither can this pattern be explained in terms of individual task items making up the episode. Instead, this pattern can only be explained by taking into account some task episode related entity that came into being at the beginning of the episode eliciting maximal deactivation, and then decreased in its load as parts of the episode were executed, causing a decrease in deactivation. We suggest that this entity was the task episode related program – cognitive structures that embodied the set of higher level commands through which various neurocognitive domains were controlled and organized across time.

Such programs are created at the beginning of task episodes and manifest in the characteristic behavior seen at step 1 of task sequence executions (Rosenbaum et al., 1983; Schneider and Logan, 2006; Farooqui and Manly, 2017), and are dismantled at task episode completions eliciting characteristically widespread end activity (Fujii and Graybiel, 2003; Farooqui et al., 2012; Crittenden et al., 2015; Simony et al., 2016). This additional activity at episode completion was also evident in all of the current experiments where the last step frequently had a significantly positive activity (Figures 2, 3, 6, 8, 9 and 14), and the estimates of time-course showed additional activity elicited at episode completions (Figures 9 and 14). The magnitude and spread of the task episode completion activity tends to be related to the hierarchical level of the completed episode whereby the activity elicited by task episode completion > subtask episode completion > sub-subtask episode completion (Farooqui et al., 2012). This further evidences the hierarchical and subsuming nature of programs, whereby completion of a subtask episode dismantles the program related to it, leaving the program related to the overarching task episode intact, hence elicits less activity than task episode completion that dismantles programs at all levels.

### Program Related Deactivation

Deactivations are ubiquitously seen during external task executions and have been attributed to the shutting down of the network involved in internal cognition (e.g. mind wandering or self-referential processing; Mason et al., 2007; Spreng et al., 2009) during periods of external attention and/or working memory (Ingvar, 1979; Shulman et al., 1997; Raichle et al., 2001). We showed that such deactivations are related to the task episodic structure of the behavior being executed. They were elicited by the beginning of a construed episode and decreased along as parts of the episode get executed, even when sequential parts of the episode were identical in terms of external attention, working memory and other control requirements. Thus, step 1 elicited a deactivation while the final step elicited activation even though both were identical in their attention and control demands.

That attention, in and of itself, cannot account for these deactivation was further illustrated by the differences in results across experiments 2A and 2B. The initial attentional demands were identical between the task episodes from these experiments, but the initial deactivation was greater in magnitude for the longer task episodes of experiment 2A. This can only be explained in terms of programs. Programs assembled at the beginning of execution have elements related to the entire duration of ensuing episode, hence programs of longer episodes will have a heavier cognitive load and cause a greater deactivation than those related to shorter episodes. These experiments also ruled out the notion that time-on-task related attentional decrement caused a decrease in deactivation across the episode length. Such decrements, if present, will be a function of time already spent on the task, so should have caused identical patterns of decrease across task episodes of experiment 2B as they did across the corresponding duration during task episodes of 2A. In contrast, we found that while activity reached baseline around 40^th^ s in 2A (the expected time of completion), they reached baseline around 18^th^ s in 2B.

We propose that programs may be the primary cause of task related deactivations. Association of such deactivations with external attention and working memory seen in past (e.g. Shulman et al., 1997; Greicius et al., 2003) may have been a result of the fact that attention, maintaining task relevant information across time (working memory) or any other form of cognitive control always occur in the context of some form of extended task episode (e.g. a block of Stroop trials), and hence may themselves be instantiated through programs. The deactivation related to the program may either be a result of the configuration of neural activities of a region to maintain program related neural structures, or be an effect of cognitive load of programs being maintained in a different region, or perhaps both. In either case, task episodes may be a neurally global phenomenon that affects nearly the entire cortex, either as activations for executing the component items making up the episode or as part of the program related deactivation (see also Gonzalez-Castillo et al., 2012).

### Program Related Deactivation is Not Limited to the DMN

Task episode related deactivation was shown by a variable set of regions that was *not* involved in executing individual task items/steps of the episode. When in experiment 1 the episode consisted of rule switching trials executed with the right hand, sequential change was shown, amongst others, by the regions associated with DMN, right somatosensory and motor regions, and primary auditory cortices. In experiment 2, the task content of the episode was simpler - covert monitoring of easily visible letter sequences - and did not involve any motor response. Now the same pattern was additionally shown by all MD as well as sensory (except visual) and motor regions. Experiment 3 showed that during task episodes executed with the right hand such deactivation was present in the hand region related to the idle left hand but not in that related to the active right hand, and during the left hand executed task episodes such deactivation was present in the hand region related to the idle right hand but in that related to the active left hand. DMN and auditory regions showed this pattern during both task episodes.

Task related deactivation is not limited to a fixed set of specialized regions and can occur in any brain region that is not currently involved in controlling or processing the task content of the steps making up the episode. The primary cause of task related deactivation was therefore not the *default* character of the deactivated brain regions or their preference for internal cognition/mind wandering, although these may been the functions of *some* of the uninvolved regions. Previously, somatosensory, motor, auditory and visual regions have been seen to deactivate during task situations when they were not involved (Allison et al., 2000; Nirkko et al., 2001; Merabet et al., 2006; Hairston et al., 2008; Linke et al., 2011). Geranmayeh et al., (2014) observed that a left fronto-temporo-parietal system involved in language production deactivated during counting and decision tasks while its mirror on the right side did the opposite. The current results suggest that these deactivations may be related to the larger phenomenon of task related deactivation traditionally characterized in reference to the DMN. These results also argue against a necessary anti-correlation between the MD and DMN regions (c.f. Fox et al., 2005). Of the MD regions some in experiment 1 and all in experiment 2 showed identical patterns of deactivation as the DMN regions.

### Task Episode related Programs

Many accounts of hierarchical cognition consider the higher level entity controlling the execution of extended behavior to be an abstract, language derived hierarchical representation of task steps, which controls execution by specifying the identity and sequence of component steps akin to the way a recipe controls cooking (e.g. Miller et al., 1960; Schank and Abelson, 1977; Cooper and Shallice, 2000). However, the pattern of deactivation seen across task episodes cannot be explained in terms of such entities. This pattern was evident in task episodes where such representations were absent because the identity and sequence of steps were not known e.g. experiment 1. While task episodes in experiment 2 were related to a hierarchical, language derived cue representation, even here the decrease in deactivation cannot be explained through such representation. What is likely to be maintained in working memory is the composite cue representation ‘DAT’ and not its individual constitutive elements, hence it is difficult to claim that deactivation decreased because elements of this representation (‘D’, ‘A’ and ‘T’) were sequentially lost. Further, its difficult to argue why a decrease in working memory load will cause an increase in fronto-parietal activity. After all, such activity typically increases with increased working memory load. Most importantly, the step linked to the same ordinal position of such representation (e.g. search 2: ‘search for A’) was associated with decreased deactivation when it occurred later in the task episode compared to when it occurred earlier. If loss of elements from such representations caused decrease in deactivation then early and late versions of the same search should be associated with identical levels of activity.

Behavioral studies also argue against such notions of the higher level entity. Evidence of assembly of this entity is seen at the beginning of task episodes where the identity, sequence and number of component steps are unknown (Farooqui & Manly, 2017). The delay in step 1 RT at the beginning of task episodes is longer before task episodes with same number of component steps but longer total duration and before task episodes with greater probability of rule switches (Farooqui & Manly, 2017; Poljac et al., 2009). All of these suggesting that the higher level entity causing the step 1 delay cannot simply be a representation of component steps, instead it is related to the magnitude of control demands of the construed task episode as well as the duration of that episode, even though the specific details of when and where control will be needed was unknown.

Even where behavior has to be executed through a memorized task sequence, the higher level entity is not a mere representation of the task sequence. When participants are given a memorized list to execute (e.g. CCSS where C and S may respectively stand for color and shape decisions to be made across sequential trials; Schneider & Logan, 2006), item 1 RT is the highest, and is related to the expected number of item-level switches in the ensuing list, hence item 1 RT is longer before a list like CSCS (with three item level switches) than CCSS (with one item level switch). Crucially, these remain the case even when the same list is iteratively executed, suggesting that something *additional* is done to the representation already in working memory before every execution, and that this includes prospectively preparing for the control demands to come.

Programs may be better thought of as cognitive entities embodying commands for bringing about various control related changes across different neuro-cognitive domains across the length of the task episode. These changes would form the overarching cognitive context for the search, selection and execution of component cognitive processes and behavioral acts making up the episode. Such hierarchical instantiation of executive commands is well recognized in motor cognition (Henry and Rogers, 1960; Rosenbaum et al., 1983, 2007; Keele, S. W., Cohen, A., & Ivry, 1990). Motor actions typically consist of a sequence of smaller acts, e.g. articulating a word may consist of a sequence of phonemic articulations. Instead of individually instantiating executive commands for each of these component acts, a motor program embodying the commands for the entire sequence is instantiated in one-go. This then unfolds across time into the seamless chain of small acts making up the overall action.

While motor programs have typically been characterized in situations where behavior seems to get executed ballistically, programs need not be limited to such instances. Programs evidenced during the execution of memorized task sequences do not ballistically translate into behavior, instead they translate into the sequence of rule-related cognitive set changes through which the correct motor acts are selected in response to the stimuli (see discussion of Schneider & Logan, 2006). When a prior knowledge of the sequence of action selection rules to apply across time is present, the commands for the creation of these rule related cognitive set changes get instantiated in one-go, embodied in one program.

Likewise, in unpredictable task episodes where identity and sequence of steps are not known in advance, the program may bring about goal related control and attentional changes in cognition through which the correct next step would be *searched* for. For example, in experiment 2, the higher-level commands that instantiated the attentional search and its related changes in the brain across the 40-second long episodes are unlikely to be instantiated as independent acts every millisecond or every second. Instead, the program evidenced in that experiment is likely to have embodied the commands for instantiating relevant cognitive changes for the whole trial episode.

Every task episode requires organization of cognition across time. Various irrelevant processes and representations need to be cleared out and maintained in abeyance across time to remove competition for limited cognitive reserves, at the same time various task relevant learnings, memories, skills, dispositions, knowledge and expectancies, and the corresponding configurational changes in various perceptual, attentional, mnemonic, and motor processes have to be brought to fore (Bartlett and Bartlett, 1932; Rogers and Monsell, 1995; Logan and Gordon, 2001). The program may be the means of achieving these widespread changes across various neuro-cognitive domains and across time, e.g. in current task episodes processing related to mind-wandering, ongoing unconscious goals, task irrelevant sensory and motor processing etc. had to be relegated. At the same time, the predictiveness of the episode was to be utilized to make anticipatory changes; e.g. knowledge that responses would be right (or left handed), visual attention limited to area around fixation, along with an implicit idea of iTis, may have been used to increase preparations and attention at times when a trial was expected and decrease when iTi was expected.

Note that most of these changes occur automatically once a task episode is embarked upon. In a visuo-motor task, for example, the doer does not deliberately or independently decide to disregard ongoing auditory and somatosensory processing, neither does she make relevant changes in auditory and somatosensory cortices through independent acts, nor does she individually execute the sequence of such changes across time. All such changes ensue automatically once a task episode is embarked upon. In fact, all deliberative decisions are made in the context of subsuming goal related cognitive changes that proceed automatically once the goal has been embarked upon.

Programs are integrally linked to goals that task episodes culminate in. Rather than seeing them as merely end states to be achieved, goals may be better conceived as cognitive structures or programs geared towards reaching that end state. Indeed a wide variety of frameworks accept that goals are important for the control and execution of tasks that lead to their achievement (James, 1890; Lewin, 1926; Greenwald, 1972; Prinz, 1987; Jeannerod, 1988; Meyer and Kieras, 1997; Gollwitzer and Sheeran, 2006). For this to happen goals have to correspond to some cognitive entity that is active during the preceding task episode (James, 1890; Kruglanski and Kopetz, 2009). Programs may be seen as cognitive structures assembled at the beginning of execution that embody the vast set of commands that will bring about such goal-related changes in various cognitive domains across time, such that at each point of the episode the cognition is in the most goal-directed state achievable with available task knowledge.

In summary, whenever cognition/behavior is parsed into task episodes a number of brain regions deactivated at the beginning and then showed gradual return to the baseline as parts of the episode are executed, suggesting the presence of episode related programs that would be assembled at the beginning and dismantle piecemeal with as parts of the episode are executed. These results make three key suggestions related to task related deactivations: (1) Programs may be the cause of these deactivations. (2) These deactivations may not be limited to a fixed set of DMN regions purported to specialize in internal cognition and or mind wandering, but may be shown by any region not currently involved in executing individual components of the task episode. (3) DMN and MD regions may not be specialized to anti-correlate during task executions; instead, depending on the nature of the task episode, they may show identical patterns of deactivation.

Authors report no conflict of interest

## Footnotes

1 Reference has been deliberately anonymised

## <u>References</u>

Allison JD, Meador KJ, Loring DW, Figueroa RE, Wright JC (2000) Functional MRI cerebral activation and deactivation during finger movement. Neurology 54:135–142 Available at: http://www.ncbi.nlm.nih.gov/pubmed/10636139 [Accessed January 26, 2017].

Andrews-Hanna JR, Reidler JS, Sepulcre J, Poulin R, Buckner RL (2010) Functional-anatomic fractionation of the brain’s default network. Neuron 65:550–562 Available at: http://www.ncbi.nlm.nih.gov/pubmed/20188659 [Accessed October 26, 2017].

Bartlett SFC, Bartlett FC (1932) Remembering: A Study in Experimental and Social Psychology. Cambridge University Press. Available at: https://books.google.com/books?hl=en&lr=&id=WG5ZcHGTrm4C&pgis=1 [Accessed September 13, 2015].

Buckner RL, Andrews-Hanna JR, Schacter DL (2008) The Brain’s Default Network. Ann N Y Acad Sci 1124:1–38 Available at: http://www.ncbi.nlm.nih.gov/pubmed/18400922 [Accessed May 26, 2018].

Cooper R, Shallice T (2000) CONTENTION SCHEDULING AND THE CONTROL OF ROUTINE ACTIVITIES. Cogn Neuropsychol 17:297–338 Available at: http://www.ncbi.nlm.nih.gov/pubmed/20945185 [Accessed October 26, 2017].

Crittenden BM et al. (2015) Recruitment of the default mode network during a demanding act of executive control. Elife 4:e06481 Available at: http://www.ncbi.nlm.nih.gov/pubmed/25866927 [Accessed January 10, 2017].

Cusack R, Vicente-Grabovetsky A, Mitchell DJ, Wild CJ, Auer T, Linke AC, Peelle JE (2015) Automatic analysis (aa): efficient neuroimaging workflows and parallel processing using Matlab and XML. Front Neuroinform 8:90 Available at: http://journal.frontiersin.org/article/10.3389/fninf.2014.00090/abstract [Accessed January 27, 2017].

De Jong R (1995) The Role of Preparation in Overlapping-task Performance. Q J Exp Psychol Sect A 48:2–25 Available at: http://www.tandfonline.com/doi/abs/10.1080/14640749508401372 [Accessed September 13, 2015].

Dosenbach NUF, Visscher KM, Palmer ED, Miezin FM, Wenger KK, Kang HC, Burgund ED, Grimes AL, Schlaggar BL, Petersen SE (2006) A core system for the implementation of task sets. Neuron 50:799–812 Available at: http://www.pubmedcentral.nih.gov/articlerender.fcgi?artid=3621133&tool=pmcentrez&render_type=abstract [Accessed June 2, 2015].

Duncan J (2006) EPS Mid-Career Award 2004: Brain mechanisms of attention. Q J Exp Psychol 59:2–27 Available at: http://www.tandfonline.com/doi/abs/10.1080/17470210500260674 [Accessed October 22, 2015].

Duncan J (2013) The structure of cognition: attentional episodes in mind and brain. Neuron 80:3550 Available at: http://www.sciencedirect.com/science/article/pii/S0896627313008465 [Accessed June 1, 2015].

Farooqui AA, Manly T (2017) Cognitive Entities Underpinning Task Episodes. bioRxiv:216796 Available at: https://www.biorxiv.org/content/early/2017/11/09/216796 [Accessed December 4, 2017].

Farooqui AA, Mitchell D, Thompson R, Duncan J (2012) Hierarchical organization of cognition reflected in distributed frontoparietal activity. J Neurosci 32.

Fox MD, Snyder AZ, Barch DM, Gusnard DA, Raichle ME (2005) Transient BOLD responses at block transitions. Neuroimage 28:956–966 Available at: http://www.ncbi.nlm.nih.gov/pubmed/16043368 [Accessed February 17, 2017].

Fujii N, Graybiel AM (2003) Representation of Action Sequence Boundaries by Macaque Prefrontal Cortical Neurons. Science (80-) 301:1246–1249 Available at: http://www.ncbi.nlm.nih.gov/pubmed/12947203 [Accessed February 4, 2017].

Geranmayeh F, Wise RJS, Mehta A, Leech R (2014) Overlapping Networks Engaged during Spoken Language Production and Its Cognitive Control. J Neurosci 34:8728–8740 Available at: http://www.jneurosci.org/content/34/26/8728.abstract7etoc [Accessed June 25, 2014].

Glover GH (1999) Deconvolution of Impulse Response in Event-Related BOLD fMRI1. Neuroimage 9:416–429 Available at: http://www.sciencedirect.com/science/article/pii/S1053811998904190 [Accessed September 26, 2017].

Gollwitzer PM, Sheeran P (2006) Implementation Intentions and Goal Achievement: A Meta-analysis of Effects and Processes. Adv Exp Soc Psychol 38:69–119.

Gonzalez-Castillo J, Saad ZS, Handwerker DA, Inati SJ, Brenowitz N, Bandettini PA (2012) Whole-brain, time-locked activation with simple tasks revealed using massive averaging and model-free analysis. Proc Natl Acad Sci U S A 109:5487–5492 Available at: http://www.ncbi.nlm.nih.gov/pubmed/22431587 [Accessed October 26, 2017].

Greenwald AG (1972) On doing two things at once: Time sharing as a function of ideomotor compatibility. J Exp Psychol 94:52–57 Available at: http://psycnet.apa.orgjournals/xge/94/1/52 [Accessed September 13, 2015].

Greicius MD, Krasnow B, Reiss AL, Menon V (2003) Functional connectivity in the resting brain: a network analysis of the default mode hypothesis. Proc Natl Acad Sci U S A 100:253–258 Available at: http://www.ncbi.nlm.nih.gov/pubmed/12506194 [Accessed October 26, 2017].

Hairston WD, Hodges DA, Casanova R, Hayasaka S, Kraft R, Maldjian JA, Burdette JH (2008) Closing the mind Js eye: deactivation of visual cortex related to auditory task difficulty. Neuroreport 19:151–154 Available at: http://content.wkhealth.com/linkback/openurl7sid=WKPTLP:landingpage&an=00001756-200801220-00004 [Accessed July 25, 2017].

Henry FM, Rogers DE (1960) Increased Response Latency for Complicated Movements and A “Memory Drum” Theory of Neuromotor Reaction. Res Quarterly Am Assoc Heal Phys Educ Recreat 31:448–458 Available at: http://www.tandfonline.com/doi/abs/10.1080/10671188.1960.10762052f.VfV7YHghun0 [Accessed September 13, 2015].

Ingvar DH (1979) ‘Hyperfrontal’ distribution of the cerebral grey matter flow in resting wakefulness; on the functional anatomy of the conscious state. Acta Neurol Scand 60:12–25 Available at: http://www.ncbi.nlm.nih.gov/pubmed/495039 [Accessed October 26, 2017].

James W (1890) The Principles of Psychology, Volume 1. H. Holt. Available at: https://books.google.co.uk/books/about/The_Principles_of_Psychology.html?id=JLcAAAAA_MAAJ&pgis=1 [Accessed September 13, 2015].

Jeannerod M (1988) The neural and behavioural organization of goal-directed movements. Oxford psychology series, No. 15. Clarendon Press.

Keele, S. W., Cohen, A., & Ivry R (1990) Motor programs: Concepts and issues. In M. Jeannerod (Ed.), Attention and Performance Xiii: Motor Representation and Control. Taylor & Francis. Available at: https://books.google.com/books?id=c_0kAQAAMAAJ&pgis=1 [Accessed September 13, 2015].

Kruglanski AW, Kopetz C (2009) What is so special (and nonspecial) about goals?: A view from the cognitive perspective. In: The Psychology of Goals (Gordon B. Moskowitz HG, ed), pp 27–55.

Lewin K (1926) Vorsatz, Wille und Bedürfnis. Psychol Forsch 7:330–385 Available at: http://link.springer.com/10.1007/BF02424365 [Accessed September 13, 2015].

Linke AC, Vicente-Grabovetsky A, Cusack R (2011) Stimulus-specific suppression preserves information in auditory short-term memory. Proc Natl Acad Sci U S A 108:12961–12966 Available at: http://www.ncbi.nlm.nih.gov/pubmed/21768383 [Accessed October 26, 2017].

Logan GD, Gordon RD (2001) Executive control of visual attention in dual-task situations. Psychol Rev 108:393–434 Available at: http://www.ncbi.nlm.nih.gov/pubmed/11381835 [Accessed February 17, 2017].

Mason MF, Norton MI, Van Horn JD, Wegner DM, Grafton ST, Macrae CN (2007) Wandering minds: the default network and stimulus-independent thought. Science 315:393–395 Available at: http://www.ncbi.nlm.nih.gov/pubmed/17234951 [Accessed October 26, 2017].

Merabet LB, Swisher JD, McMains SA, Halko MA, Amedi A, Pascual-Leone A, Somers DC (2006) Combined Activation and Deactivation of Visual Cortex During Tactile Sensory Processing. J Neurophysiol 97:1633–1641 Available at: http://www.ncbi.nlm.nih.gov/pubmed/17135476 [Accessed January 26, 2017].

Meyer DE, Kieras DE (1997) A computational theory of executive cognitive processes and multiple-task performance: Part 1. Basic mechanisms. Psychol Rev 104:3–65 Available at: http://www.ncbi.nlm.nih.gov/pubmed/9009880 [Accessed December 4, 2017].

Miller GA, Galanter E, Pribram KH (1960) Plans and the structure of behavior.

Monsell S (2003) Task switching. Trends Cogn Sci (Regul Ed) 7:134–140 Available at: http://www.ncbi.nlm.nih.gov/pubmed/12639695 [Accessed December 19, 2011].

Nirkko AC, Ozdoba C, Redmond SM, Bürki M, Schroth G, Hess CW, Wiesendanger M (2001) Different Ipsilateral Representations for Distal and Proximal Movements in the Sensorimotor Cortex: Activation and Deactivation Patterns. Neuroimage 13:825–835.

Pleger B, Blankenburg F, Ruff CC, Driver J, Dolan RJ (2008) Reward facilitates tactile judgments and modulates hemodynamic responses in human primary somatosensory cortex. J Neurosci 28:8161–8168 Available at: http://www.ncbi.nlm.nih.gov/pubmed/18701678 [Accessed October 17, 2017].

Poljac E, Koch I, Bekkering H (2009) Dissociating restart cost and mixing cost in task switching. Psychol Res Psychol Forsch 73:407–416 Available at: http://link.springer.com/10.1007/s00426-008-0151-9 [Accessed March 3, 2018].

Prinz W (1987) Ideo-motor action. In: Perspectives on perception and action, ed. H. Heuer & A. F. Sanders. In: Perspectives on perception and action (Heuer H, Sanders F, eds), pp 47–76. Hillsdale, NJ: Lawrence Erlbaum Associates.

Raichle ME, MacLeod AM, Snyder AZ, Powers WJ, Gusnard DA, Shulman GL (2001) A default mode of brain function. Proc Natl Acad Sci U S A 98:676–682 Available at: http://www.ncbi.nlm.nih.gov/pubmed/11209064 [Accessed February 4, 2017].

Rogers RD, Monsell S (1995) Costs of a predictible switch between simple cognitive tasks. J Exp Psychol Gen 124:207–231 Available at: http://doi.apa.org/getdoi.cfm?doi=10.1037/0096-3445.124.2.207 [Accessed December 4, 2017].

Rosenbaum DA, Cohen RG, Jax SA, Weiss DJ, van der Wel R (2007) The problem of serial order in behavior: Lashley’s legacy. Hum Mov Sci 26:525–554 Available at: https://www.sciencedirect.com/science/article/abs/pii/S0167945707000280 [Accessed April 8, 2018].

Rosenbaum DA, Kenny SB, Derr MA (1983) Hierarchical control of rapid movement sequences. J Exp Psychol Hum Percept Perform 9:86–102 Available at: http://www.ncbi.nlm.nih.gov/pubmed/6220126 [Accessed May 16, 2016].

Schank RC, Abelson RP (1977) Scripts, plans, goals, and understandings an inquiry into human knowledge structures. L. Erlbaum Associates.

Schneider DW, Logan GD (2006) Hierarchical control of cognitive processes: Switching tasks in sequences. J Exp Psychol Gen 135:623–640 Available at: http://www.ncbi.nlm.nih.gov/pubmed/17087577 [Accessed December 4, 2017].

Schneider DW, Logan GD (2015) Chunking away task-switch costs: a test of the chunk-point hypothesis. Psychon Bull Rev 22:884–889 Available at: http://www.ncbi.nlm.nih.gov/pubmed/25214458 [Accessed September 13, 2015].

Shulman GL, Fiez JA, Corbetta M, Buckner RL, Miezin FM, Raichle ME, Petersen SE (1997) Common Blood Flow Changes across Visual Tasks: II. Decreases in Cerebral Cortex. J Cogn Neurosci 9:648–663 Available at: http://www.ncbi.nlm.nih.gov/pubmed/23965122 [Accessed October 26, 2017].

Simony E et al. (2016) Dynamic reconfiguration of the default mode network during narrative comprehension. Nat Commun 7:12141 Available at: http://www.nature.com/doifinder/10.1038/ncomms12141 [Accessed January 10, 2017].

Spreng RN, Mar RA, Kim ASN (2009) The Common Neural Basis of Autobiographical Memory, Prospection, Navigation, Theory of Mind, and the Default Mode: A Quantitative Metaanalysis. J Cogn Neurosci 21:489–510 Available at: http://www.ncbi.nlm.nih.gov/pubmed/18510452 [Accessed October 26, 2017].

